# Glucose metabolism distinguishes TE from ICM fate during mammalian embryogenesis

**DOI:** 10.1101/826875

**Authors:** Fangtao Chi, Mark S. Sharpley, Raghavendra Nagaraj, Shubhendu Sen Roy, Utpal Banerjee

**Author notes:** Equal Contribution (Fangtao Chi, Mark S. Sharpley, Raghavendra Nagaraj). Co-Corresponding Authors; Correspondence: Mark Sharpley Utpal Banerjee.

## Abstract

The mouse embryo undergoes compaction at the 8-cell stage and its transition to 16 cells generates polarity such that the outer apical cells are trophectoderm (TE) precursors and the inner cell mass (ICM) gives rise to the embryo. We report here, that this first cell fate specification event is controlled by glucose metabolism. Glucose does not fuel mitochondrial ATP (energy) generation and glycolysis is dispensable for blastocyst formation. Glucose does not help synthesize amino acids, fatty acids, and nucleobases. Instead, glucose metabolized by the hexosamine biosynthetic pathway (HBP) allows nuclear localization of YAP1, and the pentose phosphate pathway (PPP), along with sphingolipid (S1P) signaling, activates mTOR and allows translation of AP-2γ. YAP1, TEAD4 and AP-2γ physically interact to form a nuclear complex that controls TE-specific gene transcription. Glucose signaling has no role in ICM specification, but this cascade of events constituting “Developmental Metabolism” specifically controls the fate of TE cells.

## Introduction

Following fertilization, the single-cell mouse zygote undergoes three cleavages to generate eight totipotent blastomeres. The 8-cell embryo undergoes compaction, which gives rise to tightly connected cells with barely distinguishable boundaries. As this “compacted morula” transitions to the 16-cell stage, the outer cells acquire an apical and a basal surface, while the resulting inner cells are non-polar. This pattern is generated following the 8-cell stage since compaction reorients the planes of cell division such that cells exposed to the periphery divide in a manner that distinguishes its two progeny as apical (facing outward) and basal (surrounded on all sides by other cells). The first cell-specification event that separates the totipotent cells of the morula into two distinct lineages is evident at this time. The apical blastomeres exposed to the outside ultimately differentiate to form the extraembryonic trophectoderm (TE) while the cells surrounded on all sides by their neighbors remain nonpolar and largely contribute to the pluripotent inner cell mass (ICM) at the blastocyst stage (Chazaud and Yamanaka, 2016; Leung et al., 2016; Rossant, 2018; White et al., 2018).

The preimplantation embryo floats freely within the oviductal fluid, which provides nutrients essential for the cleavage stages. The nutritional requirements of the embryo are minimal and an *in vitro* culture medium with only pyruvate, lactate, and glucose as nutrients, but lacking any amino acids, fats or proteins supports normal development through the 4.5 days of preimplantation stages (Biggers et al., 1997; Nagaraj et al., 2017). Growth factors or cytokines from the local environment are not crucial for development as early embryogenesis occurs normally without any proteins added to the medium. In this manuscript, we demonstrate that these early developmental cues are instead generated by cooperative interactions between metabolite uptake and internal signaling events. Other than the need for the three exogenously provided metabolite nutrients, the embryo is self-sufficient in producing all necessary components to sustain the variety of developmental events that are completed prior to implantation. We find that this unique environment of the preimplantation embryo is conducive to metabolic and developmental strategies that are different in many respects, and similar in others, to that seen in differentiated cells, stem cells and/or cancer tissues.

A growth medium lacking pyruvate is unable to support progression beyond the 2-cell stage (Brown and Whittingham, 1991). In a previous study we showed that pyruvate is essential for initiating zygotic genome activation (ZGA) and also for the selective translocation of key mitochondrial TCA cycle proteins to the nucleus. This unusual process allows epigenetic remodeling and ZGA (Nagaraj et al., 2017). Glucose is not required for transitioning beyond the 2-cell stage. Whittingham and co-workers established a specific requirement for glucose during the compacted morula to blastocyst transition (Brown and Whittingham, 1991). Others elaborated on these studies and concluded that, at this stage of development, and not earlier, the requirement for glucose is absolute, and glucose cannot be substituted by the metabolites pyruvate and lactate that can fully support earlier stages of development (Martin and Leese, 1995). However, in an earlier study, Fridlander reported minimal oxidation of glucose in the mitochondria (Fridhandler, 1961; Fridhandler et al., 1967). Our data support, extend and unify these early observations. We investigate how the three major arms of glucose metabolism, glycolysis, pentose phosphate pathway (PPP) and hexosamine biosynthetic pathway (HBP) might control developmental signals that are relevant to this transition.

## Results

### Glucose is Essential for the Morula to Blastocyst Transition

A minimal medium (mKSOM; see Methods) or variants lacking individual components are used in this study of preimplantation embryonic development. This modified KSOM medium, which supports the development of zygotes to the blastocyst stage (Figure 1A-G), is devoid of all proteins, including BSA, and all amino acids, including glutamine. The three metabolites, pyruvate, lactate and glucose are included in appropriate proportions resembling the *in vivo* concentrations of these nutrients (Nagaraj et al., 2017).

**Figure 1.**
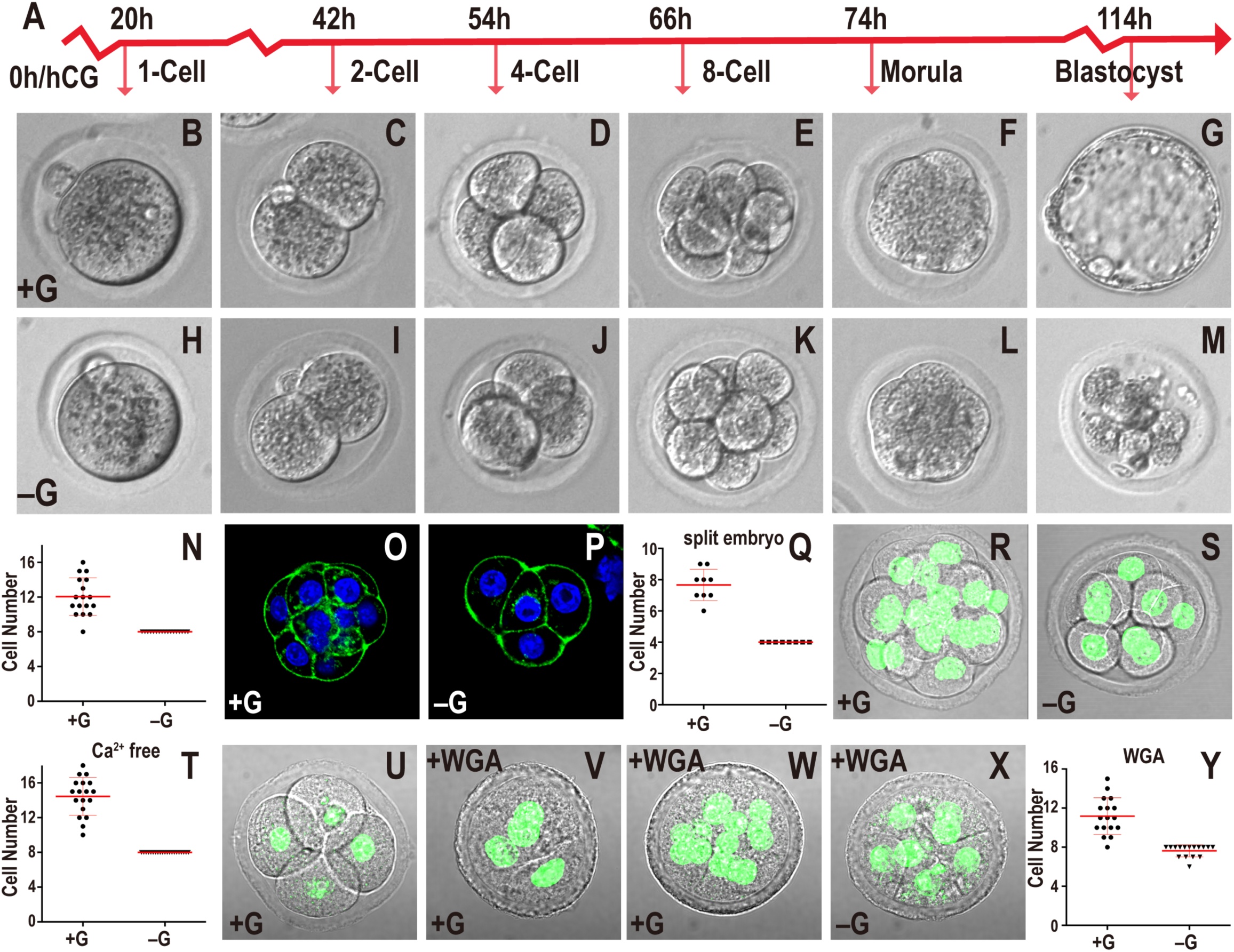
Role of glucose in early embryonic development. In this and all figures, times in hours (h) refer to time elapsed after hCG injection that induces ovulation. The normal mKSOM medium containing glucose is indicated as +G for convenience. Medium lacking glucose is marked –G. Zygotes are isolated at 18h and cultured until the specified hours (h) post hCG. All quantitative data include mean ± SD. (**A**) Timeline of developmental progression of mouse preimplantation embryos. (**B-N**) Embryos cultured in +G (**B-G**) or –G (**H-M**). The times and stages are as indicated in (**A**). In –G, the embryo fails to make a blastocyst (compare **G** and **M**), and quantitation of data at 78h shows that every embryo in –G (n=17) is blocked at the 8-cell stage (**N**). (**O-Q**) 2-cell embryos mechanically split into two individual blastomeres and grown in +G (**O**) or –G (**P**) until 78h. In both cases the embryos compact at the 4-cell stage however, quantitation (**Q**) shows that the +G split embryo is able to proceed to the 8-cell stage but the –G embryo is blocked at the 4-cell stage. Note: Since we start with a 2-cell embryo (46h) and split them, the split embryo reaches the 4-cell stage at 78h when normally an embryo is at the 8-cell stage. (**R-T**) Removal of Ca^2+^ from the medium prevents compaction in both +G (**R**) and –G (**S**) embryos shown at 78h. DAPI (green) marks nuclei. Quantitation (**T**) of data in (**R-S**) shows +G embryos contain 10-18 cells (n=18) and –G (n=21) is blocked at 8-cells although neither shows compaction. (**U-Y**) An embryo grown in +G to the 4-cell stage (56h; **U**) compacts when WGA is added (58h; **V**), and proceeds to develop beyond the 8-cell stage (n=17) (78h; **W, Y**). In the absence of glucose, the WGA treated embryos are blocked at 8-cells (n=16) (78h; **X, Y**).

In the absence of glucose, zygotes proceed normally through early cleavage stages, and undergo the compaction process (Figure 1H-M), but then block in their development precisely at the compacted 8-cell stage (Figure 1N). These arrested embryos eventually de-compact and fragment. To further investigate the temporal landmarks associated with the requirement for glucose, we mechanically split a 2-cell embryo into its single cell components and allowed each to develop. Previous work has shown that such a split embryo proceeds through the steps of preimplantation development and gives rise to a blastocyst that is smaller than normal (Casser et al., 2017). We incubate split 2-cell embryos with or without glucose and find that split embryos grown with glucose compact at the 4-cell stage, but then progress to the 8-cell stage (Figure 1O, Q) and eventually make blastocysts. Split embryos grown without glucose also undergo compaction at the 4-cell stage, however they fail to develop beyond 4 cells (Figure 1P, Q). Thus, being at the 8-cell stage is not an absolute requirement to arrest in the absence of glucose.

Next, we used techniques developed to artificially modulate the timing of compaction to investigate whether the process of compaction itself is linked to the morula block. Embryos fail to compact when grown in a Ca^2+^ free medium (Shirayoshi et al., 1983), while addition of 20 µg/ml of WGA (wheat germ agglutinin) to the medium confers compaction within 1-2 hours regardless of the stage of the embryo (Johnson, 1986; Watson and Kidder, 1988). Embryos grown in a calcium-free medium that includes glucose fail to undergo compaction but they still proceed past the 8-cell stage (Figure 1R, T). Embryos grown in medium that is both glucose and calcium free, also fail to undergo compaction, but they exhibit an 8-cell developmental block (Figure 1S, T). Finally, in the converse experiment, premature compaction by WGA, applied to a 4-cell embryo, causes rapid compaction (Figure 1U, V). In the presence of glucose, such prematurely compacted embryos continue to develop beyond the 8-cell stage, whereas without glucose, they block in development at the 8-cell stage (Figure 1W-Y). We conclude that compaction *per se* does not dictate the timing of the glucose requirement. Instead, the requirement for glucose seems to follow a counting process of an as yet unclear mechanism that operates in glucose deprived embryos. The cells could undergo many kinds of stresses under these conditions and we therefore focus here on mechanisms that operate during normal development, in the presence of glucose. More precisely, we investigate if glucose metabolism plays a role at this particular time in setting up the critical first steps that distinguish between TE and ICM lineages.

### Glucose Supports a Subset of Anabolic Processes but not Energy Production at the compacted morula stage

Based on the observation that glucose uptake increases rapidly from the 8-cell to the blastocyst stage (Leese and Barton, 1984), it is tempting to speculate (Johnson et al., 2003) that the embryo’s bioenergetic requirements at this stage are sustained by glucose. In this model, the 8-cell block could be entirely attributed to lowered energy levels in the absence of glucose. Carbon labeling using ^13^C-glucose or, for comparison, using ^13^C-pyruvate/lactate followed by mass spectrometry (MS) of the two cohorts is an unbiased approach to track the products resulting from the breakdown of these nutrients. We therefore incubated embryos throughout their developmental stages in the presence of uniformly-labeled (U-^13^C) glucose and unlabeled pyruvate/lactate. In a parallel experiment, the medium contains unlabeled (^12^C) glucose and labeled, U-^13^C pyruvate/lactate. The two metabolites, pyruvate and lactate, are labeled together since they interconvert rapidly with the carbon on lactate contributing to pyruvate (Landau and Wahren, 1992). Embryos are allowed to develop until the compacted morula stage and are then analyzed for metabolites by MS. Two different platforms are utilized to separate the metabolites. When properly optimized, the HPLC-MS method combined with the more sensitive ion chromatography system (ICS-MS) detects approximately 100 metabolites from 250 embryos per sample. Each sample is run in three biological replicates. In spite of the limitations of tissue availability, this technique enables us to map the distribution of glucose and pyruvate/lactate derived carbons with a high degree of confidence. The MS analysis identifies metabolites that derive carbons from glucose as opposed to pyruvate/lactate and also those that are not labeled with either, indicating that they are either maternally derived from the oocyte or endogenously generated from the breakdown of other stored metabolites.

Surprisingly, we find that glucose does not contribute significant levels of carbon to pyruvate as would be expected in cells with active glycolysis (Figure 2A, B). More importantly, metabolites derived from the TCA cycle are not labeled by glucose carbons either. Instead, it is exogenously derived-carbons provided by pyruvate/lactate, that solely populate TCA cycle metabolites (Figure 2B). For instance, 82% of citrate carbons are derived from pyruvate/lactate, the contribution from glucose, less than 0.5%, is close to our limit of detection. The remaining unlabeled citrate is not derived from exogenously added nutrients. Since citrate carbons are not acquired from glucose, citrate-derived fatty acids, cholesterol and several other lipids are not expected to be labeled with glucose carbons either.

**Figure 2.**
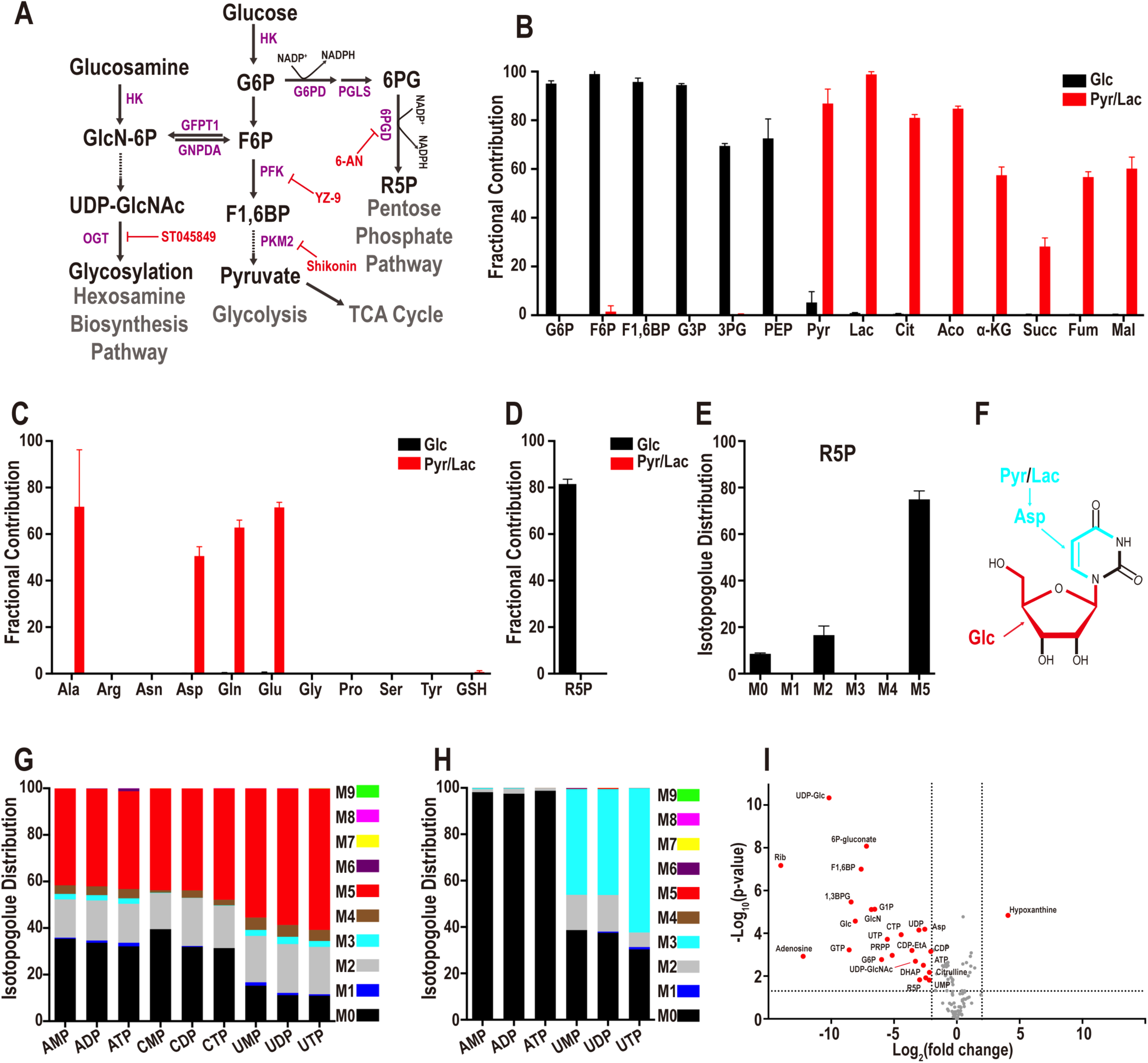
Metabolomic analysis of the compacted morula. (**A**) Representative steps in glucose metabolism. Metabolites (black), enzymes (purple) and inhibitors (red) shown are limited to those used in this work. The metabolomic analysis below is presented as mean ± SD and is obtained from 3 biological replicates, with approximately 250 embryos in each. (**B**) Embryos were cultured in either U-^13^C glucose (black bars) or, in a separate experiment, with a mixture of U-^13^C pyruvate and U-^13^C-lactate (red bars) to the 78h compacted morula stage. Metabolites were isolated and subjected to MS. U-^13^C glucose contributes to glycolytic intermediates but not to TCA cycle metabolites. U-^13^C pyruvate/lactate contributes to TCA intermediates, but not to glycolysis. (**C**) U-^13^C glucose (black) shows no detectable contribution to any of the non-essential amino acids. Contribution of U-^13^C pyruvate/lactate (red) carbons is limited to the four amino acids, Ala, Asp, Gln, and Glu. **(D, E**) Fractional contribution of U-^13^C glucose (black) and U-^13^C pyruvate/lactate (red) (**D**) to the PPP metabolite ribose-5-phosphate (R5P), and glucose isotopologue distribution of R5P (**E**). The most abundant isotopologue is M5 in which 5 of the donated carbons are from exogenous U-^13^C-glucose (black). M0 peak corresponds to unlabeled metabolites from internal resources. The origin of the M2 peak is unknown. (**F**) Structural representation of uridine. Glucose contributes 5 carbons to the ribose sugar (red), while pyruvate/lactate contributes 3 carbons to the base (light blue), via aspartate for *de novo* pyrimidine biosynthesis. (**G**) U-^13^C glucose contributes carbons to all nucleotides. No isotopologues greater than M5 are detected suggesting that only the ribose ring of the nucleotides is synthesized from glucose. U-^13^C pyruvate/lactate **(H)** contributes 3 carbons (light blue) only to pyrimidine based nucleotides (e.g. UMP, UDP, UTP), and no carbons (M0) from pyruvate/lactate are used to populate purine nucleotides (e.g. AMP, ADP, ATP). (**I**) Identification of glucose sensitive metabolites. Metabolite levels in embryos cultured in +G or –G medium were identified and quantified using HPLC-MS and ICS-MS platforms to separate the metabolites. In cases where metabolites were identified in both platforms, only the ICS data is included. 25 metabolites (red) differ by more than 4-fold and have a p-value < 0.05. Metabolites that do not meet this criterion are marked in grey.

The TCA cycle metabolite alpha-ketoglutarate (α-KG) shows a labeling pattern that is similar to citrate with 57% of α-KG carbons obtained from pyruvate/lactate, but < 0.05% from glucose (Figure 2B). Pyruvate and TCA cycle intermediates, such as α-KG, are the precursors of the four abundant amino acids, alanine, glutamate, glutamine, and aspartate that are extensively labeled by carbons from pyruvate/lactate, but are not significantly labeled by carbons from glucose (Figure 2C). The rest of the amino acids are endogenous as they are not labeled by either glucose or pyruvate/lactate (Figure 2C). Similarly, neither glucose nor pyruvate/lactate contributes to glutathione, a highly abundant tripeptide derived from glutamate, cysteine, and glycine that detoxifies the cells by removing free radicals (Figure 2C). Thus, the entire pool of glutathione in the compacted morula stage continues to be maternally derived.

The labeling data show that PPP is one of the limited numbers of metabolic pathways that is sourced by glucose at this stage of development. Labeling experiments, including isotopologue analyses, provide evidence that ribose-5-phosphate (R5P) is generated from glucose, but not from pyruvate/ lactate (Figure 2D, E). R5P is generated by either the oxidative (via G6P) or the non-oxidative (via F6P) arm of the PPP. The metabolic data suggest no significant contribution of glucose to the base components of purine nucleotides such as IMP, AMP, and GMP (Figure 2F, G; Figure S2A). Pyruvate and lactate make some contribution to the carbons used to synthesize pyrimidine bases (uracil 37%, cytosine 19%), while glucose does not label any of the detectable nucleobases (uracil < 0.02%, adenine < 0.05%, cytosine < 0.01% and thymine < 0.03%) (Figure 2G, H; Figure S2A). The lack of glucose contribution to purine bases is consistent with amino acid labeling data that finds glycine and serine are neither labeled by glucose (both < 0.2%) nor by pyruvate/lactate (both < 0.3%) (Figure 2C). In systems with proven contribution of glucose carbons to serine, this amino acid is converted to glycine which in turn contributes to purine bases (Mattaini et al., 2016).

The contribution of exogenous glucose to the ribose group is extensive. For instance, 89% of the UTP in the embryo contains ribose that is derived from glucose, while pyruvate/lactate do not provide any carbon to UTP ribose (< 0.1%) (Figure 2F-H). Thus, much of the UTP, and other ribose-ring containing nucleotides acquire carbons from glucose (Figure 2F, G). Pyrimidine nucleotides contain carbons derived from pyruvate/lactate (Figure 2F, H). Given that pyruvate/lactate do not contribute to R5P, we infer that the donated carbons belong to the base and not the ribose sugar. Thus, both glucose and pyruvate/lactate contribute to nucleotide formation at this stage, but they do so by very different pathways. In addition to the PPP, glucose derived carbons are incorporated into HBP related metabolites such as GlcNAc-1P that is labeled by and U-^13^C glucose (Figure S2B). To identify the metabolic pathways that are most sensitive to glucose omission, embryos are cultured with and without glucose present in the medium. Metabolites are extracted at the compacted morula stage and analyzed by MS following separation by the two platforms described above. Twenty-four metabolites (of total 106) are highly glucose sensitive and their levels decrease by more than 4-fold in the absence of glucose (Figure 2I). Notable examples include direct or indirect derivatives of the upper arm of glycolysis (e.g. G6P), or PPP (e.g. R5P, UTP), or the HBP (e.g. glucosamine and UDP-GlcNAc). Also notable is the observation that pyruvate and lactate remain unaffected in the metabolomic experiments that are discussed above (Figure 2I).

Taken together, the glucose utilization data presented in Figure 2, establish that under normal growth conditions, the compacted morula does not use glucose to fuel mitochondrial ATP generation, or provide carbon to synthesize amino acids, fatty acids, and nucleobases. These processes are fueled by pyruvate and lactate or by endogenous sources. Importantly, at the compacted morula stage, glucose contributes to the biosynthetic pathways that stem from upper-glycolysis, such as the HBP and the PPP, and in the absence of glucose, many of the products of these pathways decrease precipitously.

### Glycolysis, PPP and HBP Play Distinct Roles in Developmental Progression

Loss of function analysis of metabolic enzymes poses a significant challenge in the context of the preimplantation embryo. Enzymes involved in glucose metabolism are usually abundant, stable, encoded by multiple genes and maternally derived. The maternal components include RNAs, but also significant amounts of their protein products. This makes it difficult to generate loss of function phenotypes using standard genetic knockout or knockdown methodology. siRNA, shRNA and morpholino assisted strategies do not eliminate sufficient amounts of maternally derived metabolic proteins when compared with the results obtained in this system for appropriate controls. This is consistent with the observation that zygotic null alleles (such as for PDH) often bypass early phenotypes (Johnson et al., 2001), and as essential genes, we expect that maternal nulls in metabolic enzymes will affect prezygotic stages (as for PDH) (Johnson et al., 2007).

Keeping these limitations of the system in mind, we use two independent strategies to generate loss of function. The simplest is to use enzyme inhibitors whose specificity and dosage have been established in multiple previous studies (see Methods). Still, off-target effects remain a concern, and therefore we use multiple inhibitors affecting the same pathway, and if available, those that operate by different mechanisms, in order to achieve the inhibition.

Critical findings from the inhibitor studies are then confirmed and established using “Trim-Away” (Clift et al., 2017), to eliminate proteins directly instead of *via* their encoding transcripts. TRIM21 is an Fc binding E3 ligase. Hence, a complex is formed between TRIM21, the antibody and the protein of interest, and this complex is degraded by the proteasome mediated pathway. Importantly, its use along with a specific antibody completely eliminates both maternally deposited and newly synthesized proteins.

#### Glycolysis and generation of energy

The inhibitor YZ9 (Figure 2A) blocks glycolysis at the level of PFK and therefore inhibits the cascade that generates pyruvate. PPP and HBP are not affected by YZ9 (Seo et al., 2011). Strikingly, continuous inhibition of glycolysis from the 1-cell stage onward has no adverse effect on progress to the blastocyst stage (Figure 3A). While counterintuitive, surprising and contradictory to past expectations (Johnson et al., 2003), this finding of glucose independence of bioenergetics is completely consistent with the ^13^C-glucose labeling results (Figure 2B). In those experiments, we find that it is exogenous pyruvate and not glucose that contributes to the TCA cycle. Consistent with these requirements, addition of YZ9 blocks blastocyst formation if the growth medium lacks pyruvate (Figure 3A). A mere lack of pyruvate with no added YZ9 has no effect on blastocyst formation (not shown). Thus, YZ9 indeed functions as a specific inhibitor of glycolysis, but its block at the level of PFK is bypassed when pyruvate feeds into the pathway further downstream and provides carbons to the TCA cycle. We also used a second inhibitor, shikonin, which selectively blocks the function of the embryonic pyruvate kinase M2 (PKM2) (Chen et al., 2011). Similar to the results with YZ9, shikonin has no phenotypic consequence for blastocyst development, but a medium lacking pyruvate with shikonin added is unable to sustain growth beyond the morula stage even when glucose is present (Figure S3A).

**Figure 3.**
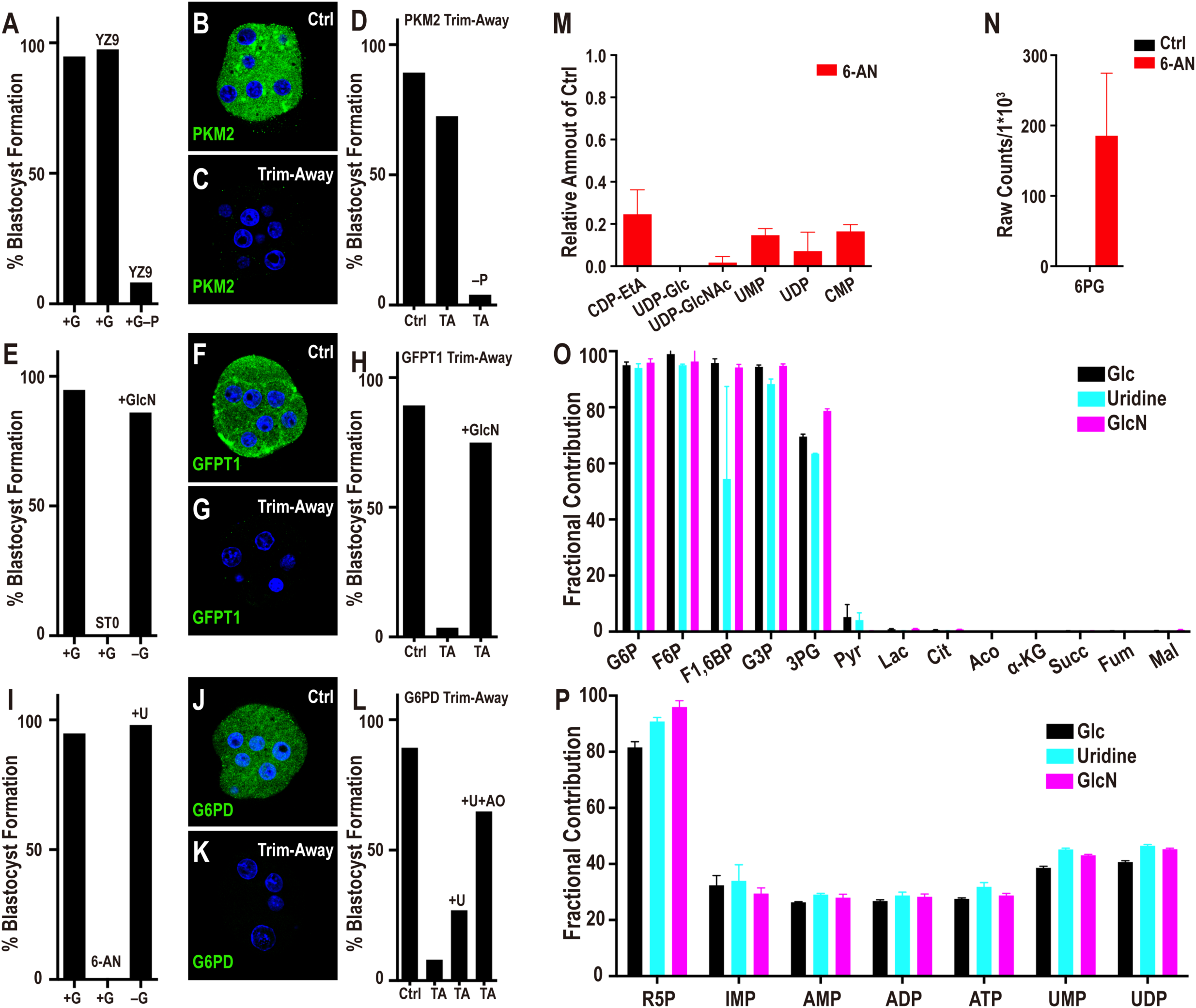
Metabolic contribution of glycolysis, PPP and HBP. (**A**) Inhibition of glycolysis (at PFK) by YZ9 (2µM) in +G has no adverse effect on blastocyst formation. However, in the absence of pyruvate (+G–P), YZ9 blocks blastocyst formation. (**B**) Control IgG antibodies injected along with mCherry-Trim21 mRNA shows robust expression of PKM2. (**C**) Injection of mCherry-Trim21 mRNA along with an antibody against PKM2 leads to complete loss of the protein. (**D**) PKM2 (lower glycolysis) depletion using Trim-Away (TA) has minimal effect on blastocyst formation. However, if the medium lacks pyruvate, blastocyst formation is no longer supported in PKM2 Trim-Away embryos. mCherry-TRIM21 channel is not shown for clarity. (**E**) Inhibition of *O*-glycosylation (HBP) by ST045849 (ST0) (0.5µM) blocks the transition from morula to blastocyst. Supplementation with glucosamine (GlcN, 1mM) fully rescues the morula block phenotype seen in –G and restores blastocyst formation. (**F**) Control IgG shows robust expression of GFPT1. (**G**) GFPT1 Trim-Away leads to complete loss of the protein. (**H**) Depletion of GFPT1 (HBP) by Trim-Away causes a morula block and prevents blastocyst formation. Glucosamine supplementation fully rescues loss of GFPT1. (**I**) Inhibition of PPP by 6-AN (2µM) blocks the transition from morula to blastocyst. Supplementation with uridine (U, 1mM) fully rescues the morula block phenotype seen in –G and restores blastocyst formation. (**J**) Control IgG shows robust expression of G6PD. (**K**) G6PD Trim-Away leads to complete loss of the protein. (**L**) Depletion of G6PD (PPP) by Trim-Away causes a morula block and prevents blastocyst formation. Uridine partially rescues G6PD loss, and this rescue is significantly better in the presence of an antioxidant (AO). Note: experiments shown in (**A, E, I**) and (**D, H, L**) were performed together, and so the respective controls are shared between the groups. (**M**) Blocking the oxidative branch of the PPP pathway with 6-AN causes a significant reduction in the levels of nucleotides and nucleotide derived metabolites. The data are normalized to the corresponding controls set at 1.0, and metabolite levels were quantified using the HPLC-MS platform. (**N**) 6-PG accumulation following 6-AN treatment. (**O, P**) Embryos were cultured in U-^13^C glucose (black), or in U-^13^C glucosamine (without glucose, light blue) or in ^13^C uridine (without glucose, purple). U-^13^C glucosamine and ^13^C uridine show a virtually identical metabolic labeling pattern as U-^13^C glucose.

Depletion of PKM2 using Trim-Away both establishes the efficacy of the method and validates the inhibitor experiment. We find that the procedure eliminates all of the detectable PKM2 protein (Figure 3B, C), and yet PKM2-depleted embryos form blastocysts with the same efficiency as in controls (Figure 3D). However, as predicted by the inhibitor analysis, in a medium lacking pyruvate, PKM2 depleted embryos are unable to transition to blastocysts (Figure 3D). Taken together, the metabolomic, inhibitor and protein loss data unambiguously establish that core glycolysis, the conversion of glucose to pyruvate, is not essential for the morula to blastocyst transition.

#### Hexosamine biosynthetic pathway (HBP)

Treatment of embryos with the commonly used inhibitors of the HBP, such as azaserine and DON that target GFPT1, results in a developmental arrest at the compacted morula stage (Figure S3A). However, these inhibitors can, under some circumstances, have off-target effects. More convincingly, we recall from the metabolomic data that UDP-GlcNAc, the critical product generated by HBP, is labeled by ^13^C-glucose and that its levels decrease in the absence of exogenously added glucose (Figure 2I). UDP-GlcNAc facilitates both *N*- and *O*-linked glycosylation. Experiments with tunicamycin that blocks *N*-glycosylation have suggested a role for this protein modification in the morula to blastocyst transition (Surani 1979). However, we later show that although *N*-glycosylation plays a role in the proper localization of apical-domain proteins, its role in cell fate determination is minor, especially in comparison to that of *O*-glycosylation in this process. ST045849 is a widely used, and very specific inhibitor of OGT (O-linked N-acetyl glucosamine transferase) function (Gross et al., 2005). When added to post ZGA 2-cell embryos, ST045849 causes a developmental arrest at the compacted morula stage (Figure 3E). The morphological phenotype of *O*-glycosylation inhibited embryos is indistinguishable from those cultured without glucose. Also, glucosamine is able to bypass and rescue the morula block caused by lack of glucose (Figure 3E) (Pantaleon et al., 2008). These data convincingly show that glucose metabolism by HBP and its role in protein *O*-glycosylation are critically important for the progression to the blastocyst stage.

To be absolutely sure of this important conclusion, we used the Trim-Away method with a specific GFPT1 antibody. This process completely eliminates the protein and also causes a compacted morula block (Figure 3F-H). Importantly, GFPT1 protein depleted embryos are rescued to form blastocysts if the growth medium is supplemented with glucosamine (Figure 3H) that directly generates glucosamine-6-phosphate (GlcN-6P), bypassing the need for an active GFPT1 enzyme.

#### Pentose phosphate pathway (PPP)

^13^C labeling by glucose demonstrates that carbons from glucose contribute extensively towards nucleotide ribose sugar biosynthesis suggesting an influx of glucose into the PPP at this stage. Furthermore, the level of nucleotides and PPP metabolites decrease substantially in embryos that are cultured without glucose. We assessed the role of PPP by inhibiting the pathway using 6-aminonicotinamide (6-AN) that is a well-established competitive inhibitor of 6-phosphogluconate dehydrogenase (PGD), the last enzyme of oxidative PPP (Downs SM, 1998; Kohler et al., 1970; Lange and Proft, 1970). With PPP thus blocked, embryos proceed normally to the 8-cell stage, undergo compaction, but arrest in development at the compacted morula stage and fail to form blastocysts (Figure 3I). The terminal product of the PPP pathway includes ribose sugars that contribute to nucleotide formation. We find that uridine added to a medium lacking glucose restores blastocyst formation (Figure 3I).

Phenotypically, 6-AN treatment closely recapitulates the developmental block caused by glucose omission. Nevertheless, 6-AN activity could, in principle, extend to other NADP^+^ dependent enzymes, although within the bounds of glucose metabolism, the PPP arm is heavily impacted. 6-phosphogluconate is an upstream metabolite of 6-AN inhibition in the PPP. We find that it is undetectable in control embryos, but accumulates to very high levels when 6-AN is added (Figure 3N).

In order to verify results from inhibitor and metabolomic analyses, we specifically depleted G6PD, the first enzyme in the oxidative branch of the PPP using Trim-Away. A monoclonal antibody that specifically recognizes G6PD is used in these experiments and we find that the G6PD protein becomes undetectable following the procedure (Figure 3J, K). Importantly, this manipulation also causes a specific developmental block at the compacted morula stage. To test the specificity of the depletion, we asked if restoring PPP activity downstream of G6PD rescues the phenotype. Partial rescue is observed when the metabolite uridine is used to supplement the culture medium of embryos with depleted G6PD (Figure 3L). This rescue is much improved when antioxidants are used along with uridine to remove the extra ROS accumulated when this enzyme function is blocked (Figure 3L).

Remarkably, of the metabolites detected in our assay, only 6 decrease substantially (more than 4-fold with P < 0.05) upon 6-AN treatment, and these metabolites all contain a nucleotide group (Figure 3M). 6-AN treatment almost completely abolishes the synthesis of nucleotide ribose. And also, uridine and cytosine derived nucleotides and their derivatives are particularly affected. The high sensitivity of UTP production to 6-AN explains why the HBP component UDP-GlcNAc is eliminated when PPP is blocked, while other metabolites belonging to the HBP pathway remain unaffected. This result highlights one of the important mechanisms of cross-talk between PPP and HBP (see below). In summary, the PPP is absolutely critical for the transition from the morula to the blastocyst stage and additionally, these data provide the first hints of a cross-talk between the different arms of the glucose metabolism pathways.

#### Cross-talk Between PPP and HBP

In the experiments described above we show that exogenously added glucosamine or uridine overcomes the morula block seen upon HBP or PPP inhibition respectively (Figure 3H, L). However, we also find that both glucosamine and uridine can independently overcome the phenotype resulting from glucose depletion (Figure 3E, I). Without glucose, both PPP and HBP are nonfunctional and it is puzzling how such an overall deficit in glucose metabolism is rescued by separately adding either glucosamine or uridine to the medium.

To investigate this question, we started with a glucose-free medium and supplemented it with either fully ^13^C-labeled glucosamine (U-^13^C-GlcN) or, in a different experiment, with uridine whose 5 ribose-ring carbons are labeled with ^13^C (^13^C-uridine). For both experiments, embryos are allowed to grow to the compacted morula stage, and metabolites are then extracted and the flow of labeled carbons to other metabolites is tracked using MS. We find that glucosamine carbons not only populate HBP metabolites (such as GlcN-6P, GlcNAc-P, UDP-GlcNAc), but interestingly, they also populate metabolites associated with the PPP (such as ribose-phosphate) and glycolytic intermediates (such as G6P) in a manner similar to that seen upon glucose labeling (Figure 3O, P). Also, similar to glucose, glucosamine does not contribute carbons to the TCA cycle, to amino acid synthesis, or to nucleobases (Figure 3O, S3B-C).

In a similar vein, labeled carbons from the ribose ring of uridine also populate metabolites in a pattern that is virtually identical to that seen when glucose is used for isotope labeling (Figure 3O, P). For instance, as with glucose, nucleotide containing metabolites, derived from the PPP, are labeled with uridine, and importantly, so are the intermediates of glycolysis and the HBP (Figure 3O, P). We conclude that uridine is metabolized and fed into glycolysis by non-oxidative PPP. As with glucose, TCA cycle intermediates, amino acids and nucleobases are not labeled by uridine (Figure 3O, Figure S3B-C). In summary, the labeling data show that either uridine or glucosamine can supply carbons to all the metabolites that G6P generates and provides evidence for extensive cross-talk between these two arms of the glucose metabolism pathways.

### Glucose Metabolism and TE Specification

The first lineage specification event in the developing embryo is later than but not far in time from when glucose becomes essential for further development leading us to investigate whether lineage specification might be causally linked to glucose utilization. We find that the canonical TE specification and maturation transcription factor CDX2 (Strumpf et al., 2005), is not expressed in embryos cultured in a glucose deprived medium or those in which either the HBP or the PPP pathway is specifically inhibited (Figure 4A-E). To our surprise, the expression of ICM markers such as OCT4 and NANOG remain entirely unaffected in the absence of glucose, and their expression also remains normal when HBP derived glycosylation or PPP is inhibited (Figure 4F-O).

**Figure 4.**
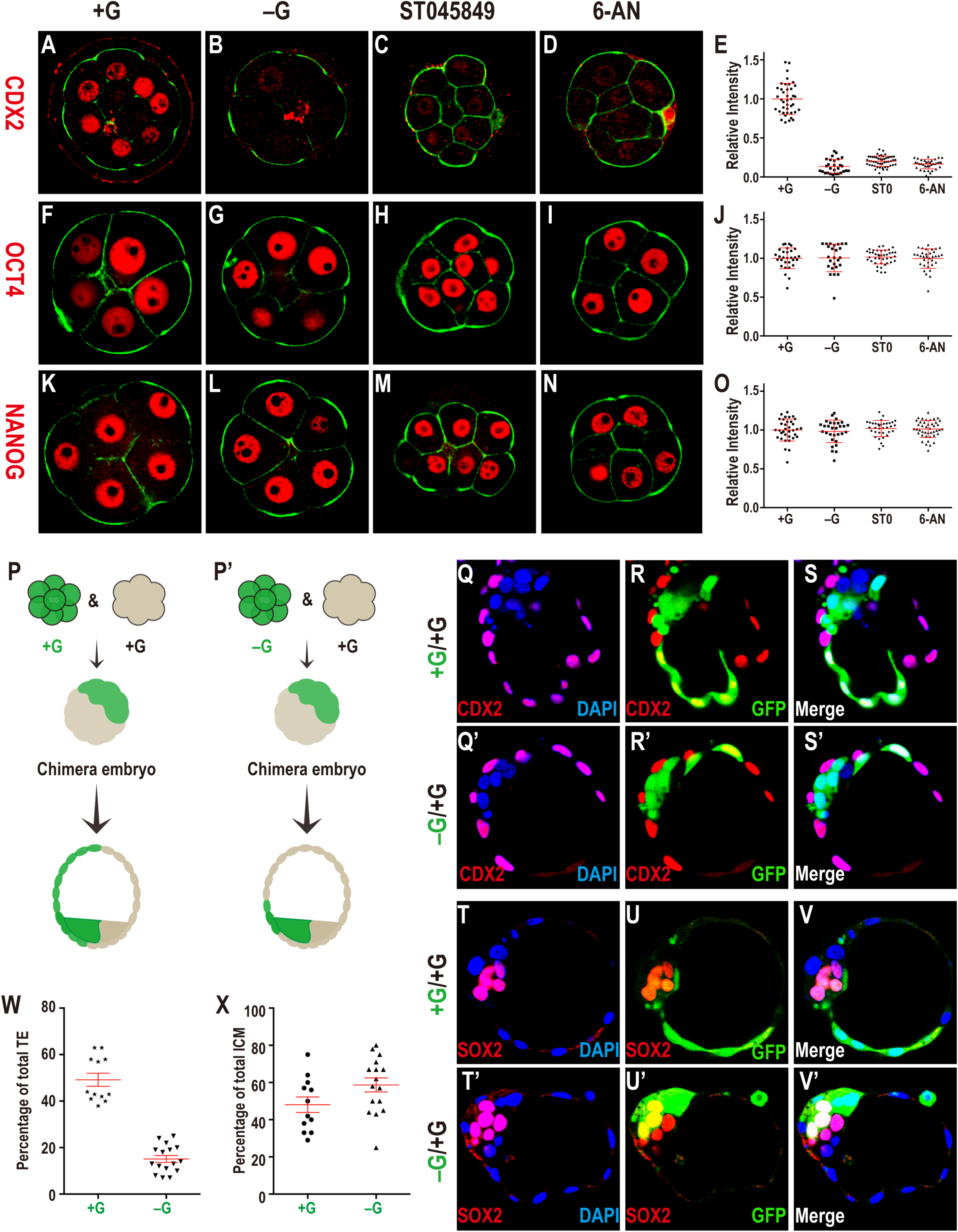
Glucose is required for TE fate initiation and specification. CDX2 (**A-E**) expression is observed in outer blastomeres in +G (**A**), but not in –G (**B**), and not when *O*-glycosylation (HBP, **C)** or PPP (**D**) is blocked. Quantitation of CDX2 data (**E**). OCT4 (**F-J**) and NANOG (**K-O**) expression in +G (**F, K**), –G (**G, L**), ST045849 (**H, M**), or 6-AN (**I, N**). Quantitation of OCT4 (**J**) and NANOG (**O**) shows that their levels are unchanged without glucose, or when PPP or *O*-linked glycosylation is blocked. (**P-X**) Chimera experiments to assess glucose function in TE specification. (**P, P’**) Schematic representation of the setup and the results, for control (**P**) and experimental (**P’**) chimeras. A GFP marked embryo (represented in green) is cultured either in +G (**P**) or –G (**P’**). An unmarked embryo (not green) is separately grown in +G to the 8-cell stage. The green and non-green embryo are fused to make a chimera that is grown without glucose (for both **P, P’**) to the blastocyst stage. In (**P**) green cells distribute equally between TE and ICM. In (**P’**) green cells rarely take on TE fate. (**Q-X**) Data supporting the scheme in (**P, P’**). (**Q-S’**) CDX2 (red, purple in merge with DAPI showing nuclei, GFP green) expression in TE cells in control (**Q-S**) and experimental (**Q’-S’**) chimeras. In the control, green cells populate both ICM and TE while in the experimental chimera, they are largely restricted to the ICM. (**T-V’**) SOX2 expression (red) is in the ICM. While green cells can be either SOX2 positive or negative in control chimera (**T-V**), in +G/–G chimera, the green cells are usually SOX2 positive (**T’-V’**) (see text). (**W, X**) Quantitation of the chimera data.

The timing of the 8-cell morula block is earlier than that of the first expression of CDX2 (16-cell stage). To determine if the results above merely represent the fact that glucose deprived embryos simply do not reach the stage that corresponds to CDX2 expression, we decided to cause a cell cycle block by an alternative method. The drug aphidicolin inhibits DNA replication. When added to early 8-cell embryos it causes a compacted morula block such that these embryos never proceed to the 16-cell stage. However, in contrast to glucose deprived embryos, 12h post-aphidicolin treatment, the 8-cell blocked embryos express normal levels of CDX2 (Figure S4A, A’). Thus, CDX2 expression does not demand that the embryo be at the 16-cell stage or later.

A morula blocked embryo never reaches the blastocyst stage which is when the fates of the TE and ICM cells become obvious, not only by markers but also by their cell position and shape. To address whether glucose is important for this later process of TE determination, rather than simply for the expression of a few isolated markers, we developed a chimera assay for our system. In these experiments, two embryos grown independently until the 8-cell stage are mechanically aggregated together to form one larger embryo that is competent to develop further to form “chimeric” blastocysts (Spindle, 1982; Tarkowski, 1961). For fate mapping purposes, one of the embryos is injected at the 1-cell stage with GFP mRNA that is efficiently translated into the green protein and marks all progeny to the blastocyst stage (Figure 4P, P’). As control, both the green and the non-green embryos are grown in normal medium containing glucose, fused at the 8-cell stage, and then further grown to the blastocyst stage in a medium lacking glucose. As expected, we find in this control that the blastomeres from the GFP and non-GFP embryos are equally competent to generate ICM and TE lineages (Figure 4Q-S, 4T-V). For the experimental group, the GFP labeled embryo is allowed to mature from the 1-cell to the early 8-cell stage in a glucose-free medium while an unlabeled embryo is grown in the normal glucose containing medium. The two embryos are aggregated and the resulting chimera is grown in a glucose-free medium until the blastocyst stage (Figure. 4Q’-S’, 4T’-V’). We find a very significant decrease (P < 0.0001) in the propensity of labeled (glucose deprived) vs the unlabeled (glucose supplemented) cells to be CDX2 positive (Figure 4Q’-S’, W). There is no significant difference in the total number of ICM or TE cells (regardless of GFP status) in chimeric blastocysts (P = 1.0). A few green cells are CDX2 positive and these are often found directly abutted to an unlabeled neighbor (Figure 4Q’-S’), and the simplest explanation is that these cells exchange glucose with the neighboring cell. The proportion of glucose deprived blastomeres in the TE and ICM are shown in Figure 4W, X. Overall, these results provide a strong basis in favor of the hypothesis that the presence of glucose is required for TE fate. The loss of TE related markers is seen at a stage earlier than the actual establishment of the fate. The chimera allow us to grow the embryo until the blastocyst stage when the fate choices are clearly discernible.

### Molecular Mechanisms Linking Glucose Metabolism to CDX2 Expression

The next step in this analysis is to probe the molecular mechanism that links CDX2 expression to PPP and HBP, since expression of CDX2 is critically dependent on the proper functioning of these two arms of glucose metabolism. The transcription factors that control CDX2 expression are well characterized in past studies (Cao et al., 2015; Rayon et al., 2014). Principal amongst these are pre-TE specification factors YAP1 and TFAP2C (also known as AP-2γ). The active YAP1 (Sasaki, 2017) is nuclearly localized within the pre-TE, but not in the ICM cells. AP-2γ is widely expressed in the 8-cell morula, but is later an important factor in controlling CDX2 in the TE (Auman et al., 2002; Cao et al., 2015; Kuckenberg et al., 2012). We focused on nuclear YAP1 and AP-2γ since they are dramatically affected in the absence of glucose (Figure 5A-F) and because these two transcription factor complexes are implicated in the direct control of CDX2 (Cao et al., 2015; Nishioka et al., 2009). Other transcription factors such as GATA3 and SBNO1 have also been linked to TE fate initiation (Home et al., 2009; Ralston et al., 2010; Watanabe et al., 2017), but since they do not show glucose dependent expression (Figure S5A-D), we have not studied these further. The expression of AP-2γ and YAP1 remains high in embryos that are arrested at the 8-cell stage following treatment with aphidicolin, and, thus, their expression is not affected by cell cycle arrest at this stage (Figure S4B-C’).

**Figure 5.**
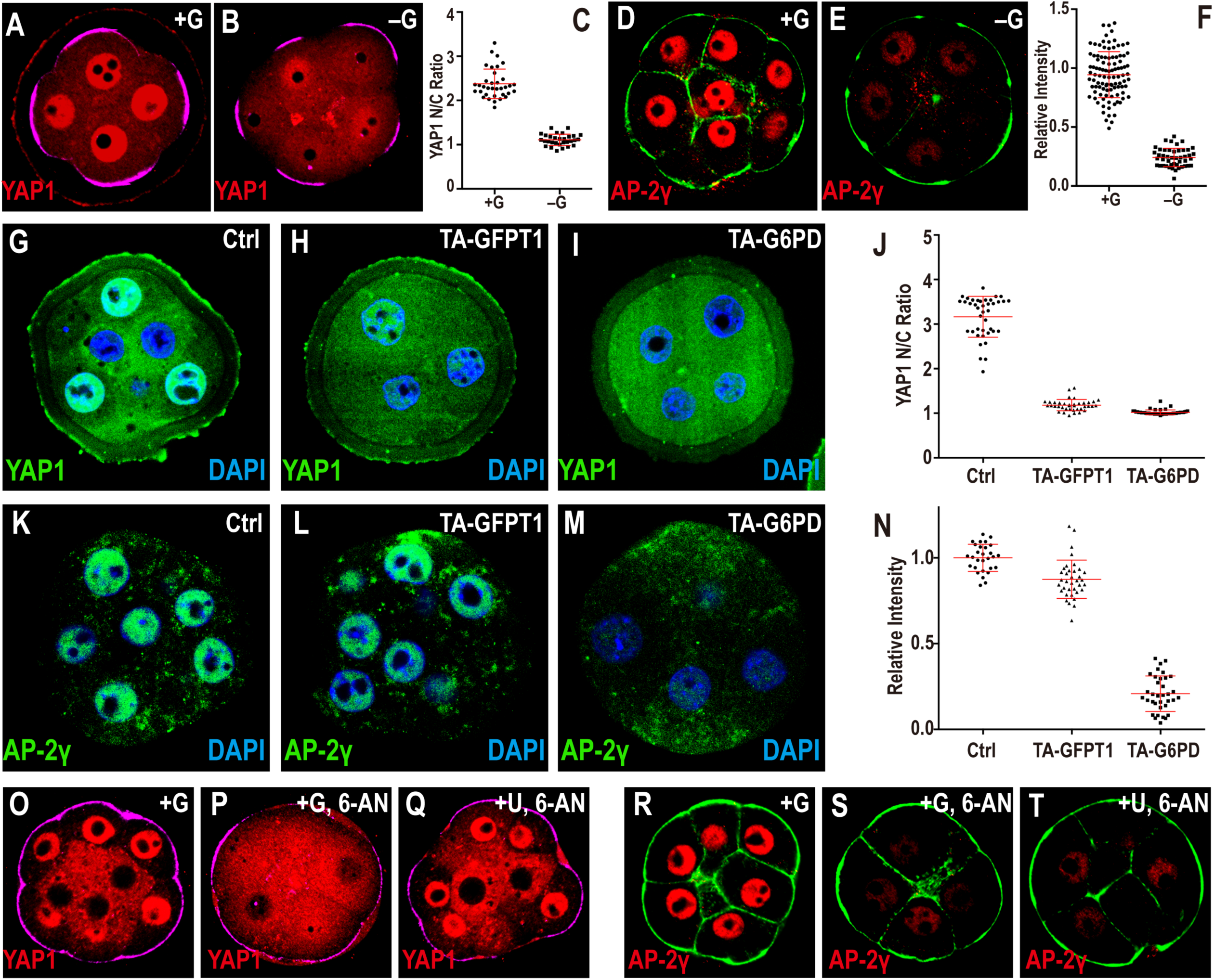
Glucose dependent activation of transcription factors. Cell membranes (green) are marked using phalloidin-FITC in (**D, E, R, S, T**), apical surfaces (purple) are marked by phospho-Ezrin/Radixin/Moesin (pERM) in (**A, B, O, P, Q**). (**A-C**) Glucose dependent YAP1 nuclear localization. In +G control (**A**) YAP1 is localized to the nucleus in pERM (purple) marked polar cells. This nuclear localization is significantly reduced in –G (**B**). (**C**) Quantitation of YAP1 nuclear to cytoplasmic (N/C) ratio. (**D-F**) In control media AP-2γ (red) is expressed in the nuclei of all blastomeres (**D**). This expression is lost in –G (**E**) Quantitation shown in (**F**). Inhibition of PPP or HBP both give results similar to that in –G (see Figure. S5E-J) (**G-N**) Loss of function using Trim-Away (TA). All embryos are injected with mCherry-Trim21 as zygotes and at the 2-cell stage with indicated antibodies. (**G**) Control, morula injected with control IgG antibody. Strong nuclear YAP1 staining (green) is seen. Trim-Away (TA) with either anti-GFPT1 (HBP) (**H**) or anti-G6PD (PPP) (**I**) results in a significant reduction in nuclear YAP1. (**J**) Quantitation of YAP1 nuclear to cytoplasmic (N/C) ratio. (**K**) Control, morula injected with non-specific IgG antibody shows robust nuclear AP-2γ (green). Injection with anti-G6PD (**M**) causes a significant reduction in AP-2γ expression while injection with anti-GFPT1 (**L**) does not influence AP-2γ expression. (**N**) Quantitation of nuclear AP-2γ. The Trim-Away data, including the effect of HBP on nuclear YAP1 localization but not on AP-2γ expression is also reflected in inhibitor data (see Figure. S5E-J). (**O-T**) Rescue experiments. YAP1 nuclear localization seen in control embryos (**O**) and lost in PPP blocked embryos (**P**) is rescued by addition of uridine (**Q**). In contrast, AP-2γ expression (**R**), also lost upon PPP inhibition (**S**) is not rescued by uridine (**T**).

Loss of HBP and PPP manifest themselves in distinct ways in affecting YAP1 vs AP-2γ. For blocking either pathway, once again, we used the reliable and specific Trim-Away technique. Loss of GFPT1 protein (HBP) using this method blocks nuclear localization of YAP1 (Figure 5G, H) but has no effect on AP-2γ expression (Figure 5K, L). Whereas, G6PD (PPP) loss by Trim-Away causes complete elimination of AP-2γ expression and YAP1 nuclear localization is also affected (Figure 5I, M). We conclude that PPP plays an important role in causing AP-2γ expression, while HBP does not (Figure 5K-M). YAP1 nuclear localization is affected when either HBP or PPP is perturbed (Figure 5G-I). These data are quantified in (Figure 5J, N).

The nuclear localization of YAP1 is severely compromised in embryos in which *O*-glycosylation is inhibited (Figure S5E, G). Recent studies have shown that nuclear localization of YAP1 requires *O*-linked glycosylation at 1-4 specific sites on the protein (Peng et al., 2017; Zhang et al., 2017). This is a likely explanation for loss of YAP1 localization when this protein modification is blocked. However, with technology currently available to us, combined with the limited availability of tissue, we cannot rule out the possibility that loss of *O*-glycosylation affects an unrelated protein that in turn affects YAP1 localization. In either case, it remains clear that HBP related *O*-glycosylation is essential for producing active nuclear YAP1 which is necessary for CDX2 expression. Phosphorylation at serine-127 also correlates with YAP1 activity (Zhao et al., 2007) but is not altered in response to glucose in our assay (Figure 6E). Consistent with the Trim-Away data, expression of AP-2γ is not at all altered when HBP-related glycosylation is inhibited (Figure S5H, J).

**Figure 6.**
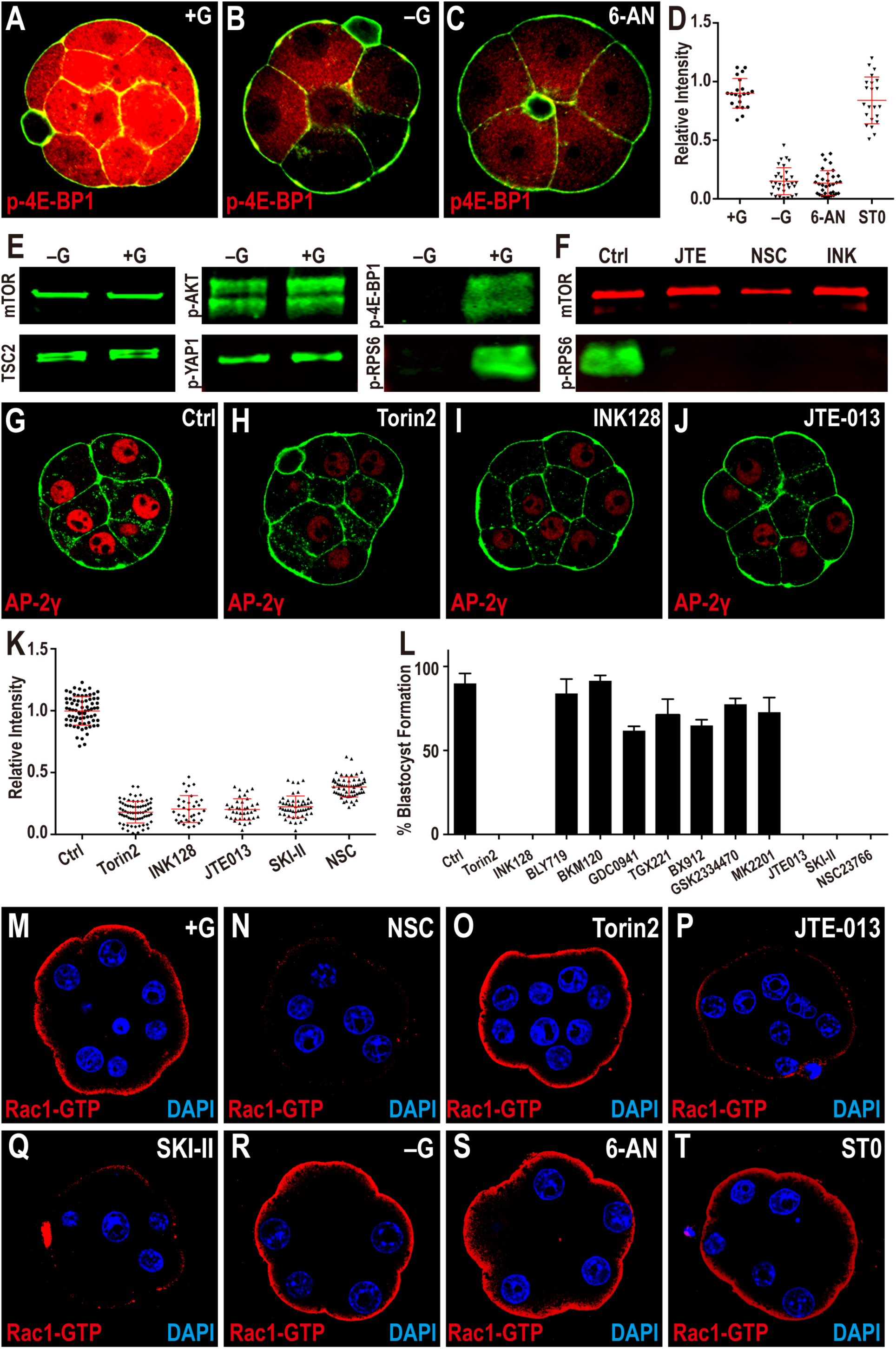
Modulation of AP-2γ Translation. (**A-D**) AP-2γ translation is controlled by PPP function. Phosphorylation of threonine 37/46 on 4E-BP1 (p-4E-BP1), a target of mTOR is expressed at high levels in the 8-cell embryo (**A**). This expression is lost in –G (**B**) or when PPP is inhibited (**C**). Quantitation in (**D**) also shows that blocking HBP does not affect p-4E-BP1. (**E, F**) Western blot analysis of members of mTOR signaling pathway and its activation. (**E**) Western blot analysis shows that total mTOR, TSC2, phosphorylation level of serine 473 on AKT1/2 (p-AKT) and serine 127 on YAP1 (p-YAP1) all remain unchanged in embryos grown without glucose when compared to +G control. In contrast, phosphorylated mTOR targets 4E-BP1 and RPS6 (p-RPS6) levels are significantly reduced in –G embryos. (**F**) p-RPS6 is eliminated when S1PR_2_ (JTE), Rac1 (NSC) or mTOR (INK) is inhibited. (**G-J**) AP-2γ expression requires mTOR and S1P signaling. The expression of AP-2γ (**G**) is eliminated in the presence of mTOR inhibitors, Torin2 and INK (**H, I**) and S1PR_2_ inhibitor JTE (**J**). An identical effect is seen on CDX2 expression (Figure. S6E-H). (**K**) Quantitation of AP-2γ expression in embryos in which the mTOR and S1P pathways are inhibited. (**L**) Quantitation shows that the morula to blastocyst transition is acutely sensitive to mTOR inhibitors (Torin2, INK128), but is not affected by any of the seven PI3K inhibitors tested (BLY719, BKM120, GDC0941, TGX221, BX912, GSK2334470, MK2201). However, blastocyst formation is extremely sensitive to inhibitors of the S1P pathway (JTE013, SKI-II) and its downstream component Rac1 (NSC23766). (**M-T**) Role of glucose and S1P signaling in controlling Rac1 expression. In control, Rac1-GTP staining is observed only on the apical surface of the polar cells (**M**). Inhibition of Rac1 (**N**), S1P receptor (**P**), and S1P biosynthesis (**Q**) cause a significant reduction in Rac1-GTP staining. In contrast, inhibition of mTOR (**O**), lack of glucose during culture (**R**), or inhibition of specific arms of glucose metabolism (PPP and HBP) (**S-T**) has no effect on Rac1-GTP levels or localization, suggesting a S1P dependent and glucose independent requirement for apical Rac1-GTP localization and mTOR activation.

Tunicamycin, a drug that blocks the HBP-related process of *N*-linked glycosylation affects the localization of apical membrane proteins (Olden et al., 1979) such as PARD6B (Figure S5K, L). However, this perturbation has no effect on either YAP1 nuclear localization or AP-2γ expression (Figure S5K-N) suggesting parallel and perhaps a more minor contribution of *N*-glycosylation compared to its *O*-glycosylation counterpart in the process of TE fate determination. Finally, it is interesting that YAP1 nuclear localization is affected by PPP. Our metabolomic analysis shows a high sensitivity of UDP-GlcNAc levels to PPP inhibition (Figure 3B). UTP is one of the most strongly affected metabolites upon PPP loss and is critically important for the generation of UDP-GlcNAc. We propose that the link between PPP and *O*-glycosylation is through this pathway’s ability to generate precursors for UDP-GlcNAc. We are able to demonstrate this to be the likely scenario by supplementing PPP inhibited embryos with uridine, a source of ribose that bypasses the PPP block to replenish UTP levels. Addition of uridine to PPP inhibited embryos fully restores YAP1 nuclear localization (Figure 5O-Q), but not AP-2γ expression (Figure 5R-T). This further demonstrates the independent regulation of AP-2γ and YAP1. Finally, although we are not yet able to measure the level of *O*-glycosylation on any individual protein, we can demonstrate that bulk *O*-glycosylation levels in the embryo are not only sensitive to HBP, but also to PPP (Figure S5O-R). This supports our hypothesis that UDP-GlcNAc generation is critically dependent on the PPP.

We next investigated the mechanism for the control of AP-2γ by PPP. From ongoing RNA-seq profiling across developmental stages we find that AP-2γ mRNA level reaches a relatively high plateau that remains virtually unchanged between the 4-cell and blastocyst stages (not shown). Importantly, this mRNA expression pattern is not dependent on the presence of glucose in the medium. In contrast, the AP-2γ protein, barely detectable until the early 8-cell stage, rises abruptly and dramatically at the compacted morula stage (Figure S5S-V). Also, the AP-2γ protein is eliminated upon deprivation of glucose (Figure 5D-F). These results first suggested to us that the control of AP-2γ expression is likely to be post-transcriptional. To test this hypothesis, early 8-cell embryos were treated with α-amanitin to inhibit transcription or with cycloheximide to inhibit translation. We find that AP-2γ protein expression remains unaffected in the presence of α-amanitin but is eliminated by cycloheximide (Figure S5W-Z). This suggests that glucose metabolism is involved in translational control of AP-2γ at the compacted morula stage.

In search of a mechanism, we turned to an analysis of the mTOR pathway that regulates protein translation in response to nutrients (Kim and Guan, 2019). To determine if mTOR activity is sensitive to glucose availability, embryos are cultured in a glucose deprived medium from the zygote stage and the level of several mTOR targets assayed in the compacted morulae. In addition to immunofluorescence analysis, we also used western blots against several components of the mTOR pathway. A key phosphorylation target of mTOR is 4E-BP1, which enables CAP-dependent translation (Sonenberg and Hinnebusch, 2009). In embryos cultured without glucose, phospho-4E-BP1 expression is abolished (Figure 6A-E) as is a second target, phospho-ULK1 (Kim et al., 2011b) (Figure S6A, B). Finally, another critical target in the mTOR-related translational cascade is the ribosomal protein S6 (RPS6), and we find that its phosphorylation level is also highly glucose sensitive (Figure 6E, S6C, D). The sensitivity of mTOR activity to glucose is not related to HBP and is traced entirely to the PPP arm of glucose metabolism (Figure 6D).

Four independent mTOR inhibitors, Torin2 and INK128 (Figure 6L), as well as rapamycin and PP242 (not shown) cause a block in the transition from the compacted morula to the blastocyst (Figure 6L). Importantly, these mTOR inhibitors abolish AP-2γ (Figure 6G-I, K) and CDX2 (Figure S6E-G) expression, but OCT4 (Figure S6I-K) and YAP1 (Figure S6M-O) remain unaffected. The fact that YAP1 nuclear localization is not mTOR dependent lends further support to the idea that the PPP dependence of YAP1 is secondarily related to its role in UDP-GlcNAc generation. Metabolized through PPP, glucose likely generates an as yet unidentified metabolite that activates mTOR and thereby translation of AP-2γ. Such a metabolite would represent a nutritional signal for mTOR activation. In this context, we note that amino acids that are typically associated with mTOR activation, or the metabolite SAM, are not glucose dependent (Figure 2I) and also that the PI3K/RTK/AKT dependent regulation of mTOR is not involved in this pathway at the 8-cell stage (Figure 6L).

### Activation of mTOR by a Signaling Lipid

A non-RTK driven mechanism for mTOR activation has been described and it involves G-protein coupled receptors (GPCRs) (Puertollano, 2019; Wauson et al., 2013). Expression analysis reveals that a small but specific class of GPCRs that bind the signaling lipid sphingosine-1-phosphate (S1P) is highly represented in the transcriptome across preimplantation stages. These attracted our attention since S1P receptors are involved in mTOR activation (Liu et al., 2009; Liu et al., 2010; Maeurer et al., 2009; Taniguchi et al., 2012). Of the several S1P receptors expressed in the embryo, the receptor S1PR_2_ follows a profile that reaches peak expression level at the morula stage.

S1P binds the extracellular domain of S1PR_2_ and activates its intracellular signaling cascade (Pyne and Pyne, 2010). The compound JTE-013, is a specific antagonist that exclusively prevents S1P binding to S1PR_2_ (Ohmori et al., 2003; Osada et al., 2002; Parrill et al., 2004). When included in the culture medium at the late 2-cell stage and beyond, JTE-013 causes a compacted morula block, and a marked decrease in phospho-4E-BP1 (Figure S6Q) and phospho-RPS6 levels (Figure 6F). Moreover, S1PR_2_ inhibition causes loss of AP-2γ (Figure 6J, K) and CDX2 expression (Figure S6H), without affecting ICM markers (OCT4 shown in Figure S6L), or the nuclear localization of YAP1 (Figure S6P). We conclude that in addition to being downstream of the PPP, the mTOR/4E-BP1/RPS6/AP-2γ cascade is also downstream of the S1P/S1PR_2_ pathway.

Although S1P binds the extracellular domain of the receptor, no lipid is exogenously provided in the medium to initiate such a signal. We therefore reasoned that S1P is synthesized within the embryo, and locally transported out of the cell. Once extruded, this lipid is able to bind the cell-surface receptor. Beyond a threshold ligand concentration, this complex would engage an intracellular signal. Several genes involved in salvage or *de novo* synthesis of sphingosine as well as sphingosine kinase (SPHK that phosphorylates sphingosine to form S1P) are all very well represented in the transcriptome at this stage of development. Inhibition of SPHK causes phenotypes that are identical to those seen upon loss of S1PR_2_ and includes a morula block (Figure 6L), loss of p-4E-BP1 (Figure S6Q), and AP-2γ (Figure 6K). Thus, like glucose, the sphingolipid pathway also regulates mTOR to control AP-2γ.

The molecular details of the link between GPCRs and mTOR are not fully understood, although several past studies have placed Rac1 as an actin cytoskeleton based downstream target of S1P/S1PRs (Gonzalez et al., 2006; Kim et al., 2011a; Reinhard et al., 2017; Zhao et al., 2009). Furthermore, a biochemical analysis has proposed that Rac1 activates mTOR by altering its localization within the cell (Saci et al., 2011). The exact activation mechanism is not the central goal of our study and we focus here instead on testing whether Rac1 is a possible link between lipid signaling and mTOR activation in our system. Following our standard assays we determined that inactivation of Rac1 function (using the Rac1-GEF inhibitor NSC23766) leads to loss of p-RPS6 (Figure 6F), p-4E-BP1 (Figure S6Q), and AP-2γ (Figure 6K) and a failure to make the compact morula to blastocyst transition (Figure 6L).

The active, GTP bound form of Rac1 is detected at the apical surface of the 8-cell embryo cultured in the presence of glucose (Figure 6M) and as a control we demonstrate that Rac1-GTP staining is lost when the interaction between Rac1 and its GEF is inhibited (Gao et al., 2004) (Figure 6N). Rac1-GTP localization is unchanged when mTOR is inhibited (Figure 6O). Importantly however, expression of Rac1-GTP is completely lost when either S1PR_2_ or SPHK is inhibited (Figure 6P, Q). These results place Rac1 upstream of mTOR and as a downstream target of S1P/S1PR_2_.

Interestingly, in sharp contrast to the dependence of Rac1 on lipid signaling, absence of glucose, or inhibition of either PPP or HBP has no effect on Rac1-GTP expression (Figure 6R-T). In other words, Rac1-GTP expression is S1P signaling dependent and glucose signaling independent. Yet, loss of either S1P or glucose (PPP) generated signal causes loss of mTOR function and prevents mTOR-induced downstream translational control. We conclude that S1P/S1PR_2_/Rac1 and glucose/PPP provide parallel inputs into mTOR. Dual inputs usually regulate mTOR (Gonzalez and Hall, 2017). For the 8-cell embryo, these are the signals generated by a lipid and an unidentified glucose (PPP) metabolism byproduct.

### Functional Integration of HBP and PPP Inputs through YAP1/TEAD4 and AP-2γ

The TE specific marker CDX2 is downstream of YAP1 and AP-2γ and published literature has shown that these two proteins participate in the direct transcriptional activation of CDX2. Consistent with these results, the proposed promoter and enhancer regions of CDX2 contain both TEAD4 and AP-2γ binding sites (Cao et al., 2015; Rayon et al., 2014). TEAD4 is the DNA binding component of the YAP1/TEAD4 transcription complex (Li et al., 2010), and we wondered if AP-2γ functions separately or as part of a larger complex with YAP1/TEAD4. This question was addressed using co-immunoprecipitation assays. Due to the very limited availability of embryos, these experiments are performed in extracts of HEK293T cells that are transfected with and express uniquely tagged YAP1, AP-2γ and TEAD4 proteins. The results from the co-immunoprecipitation assays unambiguously support a physical association between YAP1/TEAD4 and AP-2γ that would result in the formation of a larger transcription complex (Figure 7A-C).

**Figure 7.**
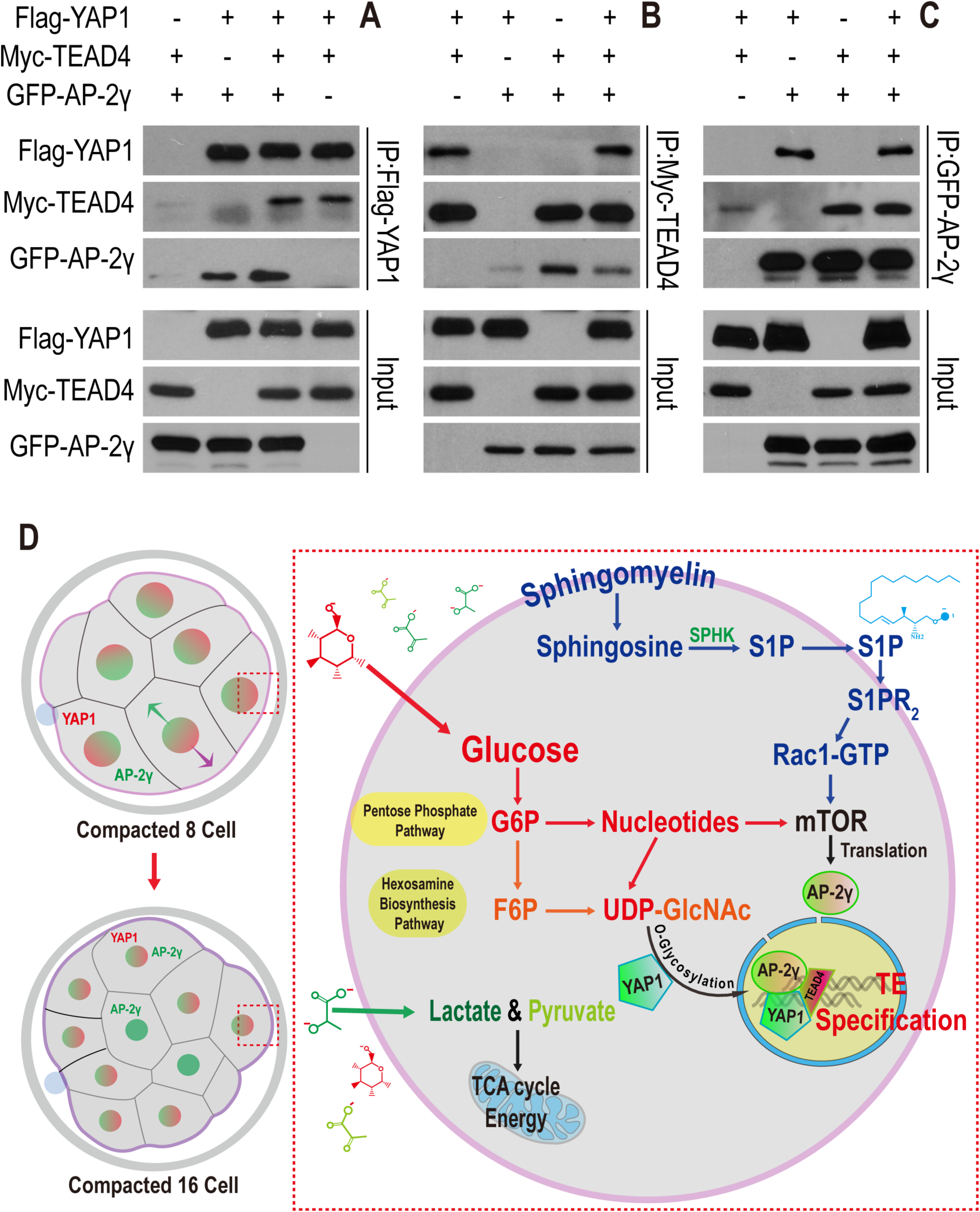
YAP1/TEAD4 and AP-2γ form a complex. (**A-C**) Flag-YAP1, Myc-TEAD4, and GFP-AP-2γ are over-expressed in HEK293T cells and cell lysates subjected to immunoprecipitation (IP) using specific antibodies (anti-Flag, anti-Myc, and anti-GFP) as indicated. Anti-Flag antibody immunoprecipitates YAP1 (as expected) but also co-immunoprecipitates AP-2γ and TEAD4 (**A**). Likewise, immunoprecipitation with anti-Myc also results in the co-precipitation of AP-2γ and YAP1 (**B**) and anti-GFP co-precipitates both YAP1 and TEAD4 (**C**). (**D**) A model for metabolic control of preimplantation development by glucose.

Based on all of the evidence presented in this paper, we conclude that in the 8-cell embryo, AP-2γ translation is controlled via the PPP arm of glucose metabolism, and by lipid signaling. The next cell division generates apical and basal cells (Figure 7D), and at this point, YAP1 is active (by glucose independent mechanisms (Sasaki, 2017)) only in apical cells. However, its nuclear localization is dependent on the glycosylation function of the HBP arm of glucose metabolism. Thus, YAP1 and AP-2γ are found together in the nucleus of the outer cells in a restricted pattern that precludes the future ICM. The outer cells initiate CDX2 expression and later give rise to the TE. This is a remarkable example of different aspects of metabolism ultimately controlling the formation of a transcriptional complex that helps discriminate between the two earliest lineages that will ultimately distinguish the embryonic from extraembryonic tissues during mammalian development (Figure 7D).

## Discussion

In this study, we have used the mouse preimplantation embryo as a model to investigate the fundamentally important, but relatively understudied relationship between signaling and metabolic pathways during normal development. The interplay between metabolic pathways and proliferation or differentiation has been studied extensively in the context of cancer cell expansion and tumor growth (Pavlova and Thompson, 2016). Although some *in vivo* assays for cancer metabolism are now available (Kang et al., 2018), they are not easy to analyze and a vast majority of such studies are done in transformed cell-lines with heterogeneous and often unknown genetic background. Ironically, metabolic control of normal development is not any easier to study because it is difficult to extract the metabolic influence on signaling pathways.

These properties are closely intertwined and work in concert to generate cellular form and function. Furthermore, most metabolic cascades are indispensable for cell viability, and phenotypic analysis of total loss of its components is often achievable in only a select few non-mammalian genetic model systems. Thus, the field of mammalian “Developmental Metabolism” presents many interesting challenges for future studies.

Several unique features of metabolic control make the preimplantation mammalian embryo a superb system in which to study metabolic control of normal development. The solitude of the mammalian embryo floating about in a small volume of oviductal fluid necessitates that the embryo be self-sufficient and require minimal input from its environment. In fact, in our modified IVF-inspired medium, lacking all but three nutrients and salts, signal transduction activities that require cell-environment interactions are essentially non-existent. This allows easier access to metabolic pathways that control all major cellular events in the preimplantation embryo.

Amongst the metabolic curiosities of the system, we find that glucose and glycolysis do not function to generate energy since glucose does not fuel the TCA cycle and does not support anabolic pathways that are TCA cycle dependent. Pyruvate takes over these functions and the prominence of the lactate/pyruvate system over glucose for fulfilling bioenergetic needs is a metabolic signature of the preimplantation embryo. We expect that this metabolic landscape would be quite different once the embryo implants and is in contact with the mother’s vasculature. Cancer cells usually rely on aerobic glycolysis for their high energy needs, although when studied *in vivo*, cancer cells are known to utilize pyruvate/lactate to fuel their mitochondria (Faubert et al., 2017; Hui et al., 2017).

In the embryo, fatty acids and cholesterol, which are generated via citrate, are not synthesized from glucose, which is an important precursor of these lipids in cancer cells. Similarly, in stark contrast to cancers (Locasale et al., 2011; Snell, 1984), glucose does not contribute carbons to amino acids including glycine and serine or to purine nucleobase synthesis. In cancer cells, amino acid and *de novo* purine base synthesis are often critical roles of glucose. Interestingly, in the embryo, most amino acids are not labeled by pyruvate either and are therefore made available from endogenous sources such as by degradation of proteins.

In spite of its limited role in glycolysis, glucose is absolutely required for development beyond the morula stage because of the important function of PPP that generates NADPH and ribose sugars, and also HBP that is essential for protein glycosylation. Bioenergetic needs could be met by a plethora of metabolites, for example, pyruvate in the embryo and glutamine in many cancer cells. But only glucose provides anabolic and protein modification functions that are essential for lineage specification, differentiation and morphogenesis during the morula to blastocyst transition. Keeping the flux through glycolysis at a minimum preserves glucose derivatives for the PPP and the HBP that are normally minor arms of the pathway.

In this context, one of the surprising findings of this study is the relatively free exchange of carbon units between the metabolites generated in PPP, HBP and glycolysis in spite of the divergent function of each arm. It is as yet unclear, and is worth investigating, if this extent of cross-talk occurs in other systems and whether low activity through the PFK-step of glycolysis is especially conducive to exchange of metabolite carbons between the arms.

By their very nature, metabolic pathways are expected to have a highly pleiotropic set of functions, affecting a multitude of cellular processes. In spite of its rich history, this wide range of functions has led to their unfortunate characterization as “house-keeping”. Quite to the contrary, we show here, as we have earlier shown in *Drosophila* (Mandal et al., 2005; Owusu-Ansah and Banerjee, 2009), that metabolic and signal transduction pathways are finely orchestrated in a way that allows the morphological and cell fate transitions that are critical for the generation of a complex structure.

It has to be the case that PPP and HBP control a larger swath of protein functions than has been investigated here. HBP must regulate the glycosylation of a large number of proteins, but in focusing on the specific system of interest, amongst many that could be affected, we find that HBP-dependent *O*-glycosylation, likely directly or perhaps through an intermediary is causally responsible for YAP1 nuclear localization in the apical cells in which it is activated by independent means. This process is largely independent of PPP except to the extent of PPP being required to make UTP and therefore UDP-GlcNAc that is essential for glycosylation within HBP. In a similar way, AP-2γ expression is controlled by mTOR, which in turn is activated by both lipid signaling and by an unidentified PPP generated metabolite. AP-2γ forms a complex with YAP1/TEAD4 in only those apical cells that have all three proteins in the nucleus. This activates TE-specific markers that are downstream. The HBP has no role to play in AP-2γ regulation.

The specificity of the events governed by this system is also highlighted by the fact that specification of ICM fate does not require glucose (neither HBP nor PPP) at all. The precise temporal and spatial combination of relatively pleiotropic components and mechanical forces that generate initial polarity (Nance, 2014) is able to give rise to this specificity such that the nutrient pyruvate provides energy, glucose/PPP activates mTOR and makes nucleotides, glucose/HBP allows the discrimination of a transcriptional event that takes place in apical and not in basal cells. Salvage and perhaps synthesis of signaling lipids provides a temporal indication of nutrient sufficiency. The net result is the separation of the TE lineage from ICM.

The preimplantation embryo is an excellent system to investigate the process of “Developmental Metabolism” because of the prominent role that metabolism plays during this time. In spite of lacking a large yolk seen in non-mammalian vertebrates and the limitations placed on resources due to the nature of internal fertilization and development, the mammalian embryo nevertheless is essentially self-sufficient in its resources. Its built-in metabolic mechanisms compensate for the paucity of extracellularly triggered signal transduction events. Later in development, these functions of metabolic pathways are likely preserved even though they might be masked by increased interactions between cells and their environment. One can readily appreciate that any imbalance between these two fundamentally important sets of pathways could lead to developmental disorders and malignancies.

## Acknowledgments

We would like to thank all members of our laboratory for their suggestions and support. Yonggang Zhou performed RNA seq analysis that was used in this manuscript. We thank Daniel Braas, and Johanna ten Hoeve-Scott at the UCLA Metabolomics Center for their generous help with metabolomics analysis. We appreciate many inputs by Tom Graeber, Heather Christofk and Hilary Coller. Christofk and Coller also helped by providing several antibody reagents. We thank Wei Liao, Qin An, Wanlu Liu and Steve Jacobsen for help with RNA sequencing experiments and bioinformatics analysis of RNA expression data, and Huachun Liu for help with molecular cloning and related molecular analysis. We thank Bin Gu at Sick Kids, Toronto for useful suggestions including for microinjection, and Bin Zhao at Zhejiang University for valuable discussions.

This work was supported by the NIH Director’s Pioneer Award to U.B. (DP1DK098059). Brief early support for this work was provided by MOD grant (1-FY17-788). U.B. is supported by the NCI grant R01CA217608. F.C. is supported by a China Scholarship Council Award and a California Institute for Regenerative Medicine pre-doctoral fellowship. Finally, we are grateful to Owen Witte and the Broad Stem Cell Research Center for several innovation awards and for continued support.

## Author Contributions

F.C., M.S.S., R.N., S.S.R, and U.B. conceived the project. F.C., M.S.S, and R.N. performed experiments and wrote the manuscript. S.S.R. set up the embryo culture system during the early phases of this work. U.B. supervised the project, secured funding, and wrote the manuscript.

## Experimental Model and Subject Details

### Mouse embryo culture

All animal care and procedures used in this study are approved by the Animal Regulatory Committee (ARC) of the University of California at Los Angeles (UCLA).

Mouse zygotes and preimplantation embryos were collected from super-ovulated 4-week old C57BL/6J X C3He (Jackson Labs) F1 females. Mice were super-ovulated by peritoneal injection of 7.5 IU of PMSG (Pregnant Mare Serum Gonadotropin) to stimulate egg production, followed by 7.5 IU of hCG (human Chorionic Gonadotropin) 48h after PMSG. Embryos were obtained by mating the super-ovulated females with C57BL/6 X C3He F1 males. Mating was confirmed by the presence of the vaginal plug. For isolation of fertilized 1-cell zygotes, super-ovulated females were euthanized 18h post hCG and zygotes were dissected out of the ampulla in the oviduct. The embryo cumulus complexes were treated with 300µg/ml of hyaluronidase to disperse the cumulus cells, washed in mKSOM medium without pyruvate/glucose and transferred to the appropriate culture medium (+G or –G) and cultured at 37°C in 5% CO_2_. All mouse embryos used in this study were cultured in a modified KSOM medium whose composition is identical to KSOM in salts, glucose, lactate and pyruvate (95mM NaCl, 2.5mM KCl, 0.35mM KH_2_PO_4_, 0.20mM MgSO_4_, 25mM NaHCO_3_, 1.71mM CaCl_2_, 0.01 mM EDTA, 0.20mM glucose, 10mM lactate, 0.20mM pyruvate) but was devoid of all amino acids and BSA. The medium also contained 0.01% PVA (polyvinyl alcohol).

### Method Details

#### Reagents

The following antibodies and drugs were used in this study. Fluorescein Phalloidin (Thermo Fisher #F432, at 1:2000 dilution), mouse anti-CDX2 (Biogenex #MU392A-UC), goat anti-OCT3/4 (Santa Cruz #sc-8628), rabbit anti-NANOG (ReproCELL #RCAB0002P-F), mouse anti-YAP1 (Abnova #H00010413-M01), rabbit anti-AP-2γ (Santa Cruz #sc-8977), mouse anti-AP-2γ (Santa Cruz #sc-12762), rabbit anti-phospho-Ezrin (Thr567)/Radixin (Thr564)/Moesin (Thr558) (CST #3141), rabbit anti-SBNO1 (Abcam #ab122789), rabbit anti-GATA3 (Santa Cruz #sc-9009), rabbit anti-PARD6B (Santa Cruz #sc-67393), mouse anti-O-Linked N-Acetylglucosamine [RL2] (Abcam #ab2739), rabbit anti-phospho-4E-BP1 (Thr37/46) (CST #2855), rabbit anti-phospho-S6 Ribosomal Protein (Ser235/236) (CST #4858), rabbit anti-mTOR (CST #2983), rabbit anti-TSC2 (CST #4308), rabbit anti-phospho-AKT (Ser473) (CST #4060), rabbit anti-phospho-YAP (Ser127) (CST #4911), rabbit anti-phospho-ULK1 (Ser757) (CST #6888), and mouse anti-Rac1-GTP (NewEast #26903, at 1:500 dilution). All the antibodies are used at 1:100 dilution for IF and 1:1000 dilution for WB.

Mouse IgG1 isotype control (Abcam #ab18447, 1mg/ml), rabbit IgG isotype control (Abcam #ab199376, 1mg/ml), rabbit anti-GFPT1 (Abcam #ab236053, 1mg/ml), rabbit anti-PKM2 (D78A4) (CST, in carrier-free formulation, 1mg/ml), and rabbit anti-G6PD (D5D2) (CST, in carrier-free formulation, 1mg/ml) were used for Trim-Away experiments.

Various chemical drugs were purchased as indicated: Lectin from Triticum vulgaris (Sigma #L0636, 20µg/ml), YZ9 (Cayman Chemical #15352, 1µM), ST045849 (TimTec, 0.8µM), Tunicamycin (Sigma #T7765, 5µM), 6-Aminonicotinamide (6-AN) (Sigma #A68203, 2µM), N-Acetylcysteine (NAC) (Sigma #A7250, 1mM), Hypotaurine (Sigma #H1384, 1mM), Shikonin (Cayman Chemical #14751, 1µM), Azaserine (Sigma #A4142, 1µM), DON (Sigma #D2141, 20µM), Aphidicolin (Cayman Chemical #14007, 5µM), α-Amanitin (Cayman Chemical #17898, 100µg/ml), Cycloheximide (Cayman Chemical #14126, 10µg/ml), Torin2 (Cayman Chemical #14185, 10nM), INK128 (Cayman Chemical #11811, 100nM), JTE-013 (TOCRIS #2392, 1µM), SKI-II (Echelon #B-0024, 0.8µM), NSC26722 (TOCRIS #2161, 100µM), BYL719 (Cayman Chemical #16986, 1µM), BKM120 (Cayman Chemical #11587, 0.5µM), GDC0941 (Cayman Chemical #11600, 0.5µM), TGX221 (Cayman Chemical #10007349, 0.5µM), BX912 (Cayman Chemical #14708, 100nM), GSK2334470 (Cayman Chemical #18095, 50nM), MK2206 (Cayman Chemical #11593, 200nM).

GFP mRNA (TriLink #L-7201), pGEMHE-mCherry-mTrim21 (Addgene #105522), pCMV6-YAP1 (OriGene #MR219506), pCMV6-TEAD4 (OriGene #MR219506) and pCMV6-AP-2γ (Origene #MR207174).

#### Antibody staining for immunofluorescence

Embryos were fixed in 4% paraformaldehyde for 30 min at room temperature, permeabilized for 30 min in PBS with 0.4% Triton (PBST), blocked in PBST with 3% albumin (PBSTA) for 30 min and incubated with the desired primary antibody in PBSTA overnight at 4°C. The following day the embryos were washed in PBST 4 times for 10 min each, blocked with PBSTA, incubated with the appropriate secondary antibody (1:500 dilution) and DAPI overnight at 4°C. Embryos were washed again 3 times for 10 min each in PBST, deposited on glass slides and mounted in Vectashield (Vector Laboratories) medium. Images were captured using Zeiss LSM700 or LSM880 confocal microscopes.

#### Inhibitor experiments

Most of the inhibitors were used to treat the embryos in the oil-free condition at the late 2-cell stage (at 50h post hCG). Early 4-cell stage embryos (56h post hCG) were treated with 20µg/ml WGA for 2 hours to induce premature compaction. Early 8-cell stage embryos (66h post hCG) were treated with 10µM of aphidicolin, 100µg/ml of α-amanitin, or 10µg/ml of cycloheximide for 12 hours (until 78h post hCG) in the oil-free condition. The embryos treated with different drugs were fixed in 4% paraformaldehyde for 30 min for immunofluorescence.

#### Measurement of metabolite levels

The metabolites are measured using the procedure described previously (Nagaraj et al., 2017; Sullivan et al., 2018). For samples analyzed under different conditions, embryos were very briefly washed in ice-cold 150mM ammonium acetate, and transferred into 80% methanol. Dried metabolites were re-suspended in 20µl 50% ACN and 5µl was injected for chromatographic separation on the UltiMate 3000 RSLC (Thermo Scientific) UHPLC system which is coupled to a Thermo Scientific Q Exactive that is run in polarity switching mode. For the UHPLC separation, the mobile phase comprised (A) 5mM NH_4_AcO, pH 9.9, and (B) ACN. A Luna 3mm NH2 100A (150 × 2.0mm) (Phenomenex) column was used with a 300 μL/min flow-rate. The gradient ran from 15% A to 95% A in 18 min, followed by an isocratic step for 9 min and re-equilibration for 7 min. The remainder of the sample was diluted 2.67-fold and 10µl were applied to a Thermo Scientific Ion Chromatography System (ICS) 5000 that is also coupled to a Thermo Scientific Q Exactive run in negative polarity mode. The gradient ran from 10mM to 90mM KOH over 25 min with a flow rate of 350µl/min. The settings for the HESI-II source were: S-lens 50, Sheath Gas 18, Aux Gas 4, spray heater 320°C, and spray voltage −3.2 kV. Metabolites were identified based on accurate mass (±3 ppm) and retention times of pure standards. Relative amounts of metabolites and contribution of ^13^C labeled nutrients (following correction for natural abundance) were quantified using TraceFinder 3.3. The data were analyzed using the Prism software package.

#### Trim-Away Experiment

The Trim-Away procedure was performed as previously described in the published literature with minor modifications (Clift et al., 2018). mCherry-Trim21 mRNA was *in vitro* transcribed using the HiScribe™ T7 ARCA mRNA Kit according to the manufacturer’s instructions. Specific antibodies in carrier-free solution (PBS) were purchased from CST or Abcam as listed above, with the stock concentration 2mg/ml. Then the mCherry-Trim21 mRNA (final concentration: 400ng/µl) and specific antibody (final concentration: 1mg/ml) were mixed with each other, and the detergent NP-40 so that the final concentration of NP-40 is 0.01%. The mixture is then microinjected into 1-cell (20h hCG) and 2-cell stage (46h hCG) embryos using a FemtoJet microinjector. A negative capacitance is generated using micro-ePORE (World Precision Instruments) to assist the 2-cell microinjection (Gu et al., 2018; Zernicka-Goetz et al., 1997). The developmental progression of these Trim-Away embryos was monitored until 114h post hCG or the embryos were fixed at the morula stage for further immunofluorescence analysis.

#### Aggregation of chimeric embryos

Embryos were labeled by microinjecting 100ng/µl GFP mRNA at 1-cell stage (20h hCG), and then cultured in +G or –G medium until the morula stage (76h hCG). Unlabeled embryos were cultured in +G medium until the compacted morula stage (76h hCG) as aggregation partners. The zona pellucida was removed by incubating each group of embryos in pre-warmed acidic tyrode’s solution for 30 seconds at 37°C. One of the zona-free embryos from GFP labeled (–G, or +G in the control group) and unlabeled (+G in both control and experimental) groups were immediately aggregated and cultured in small drops (10µl) of –G medium. The drops were covered with mineral oil until blastocyst stage (114h hCG). Each small drop of the –G medium contains a tiny well that is created by pressing and rotating the tip of a ball-point pen on the surface of the tissue culture dishes. The significance of the difference in the propensity to form TE and ICM cells was determined using a 2X2 contingency table that contained the number GFP labeled and unlabeled cells in each population and was analyzed using Fisher’s exact test.

#### Cell culture, transfection, immunoprecipitation

HEK293T cells were cultured in DMEM (Invitrogen) containing 10% FBS (Invitrogen) and 50 μg/ml penicillin/streptomycin (P/S). Transfection with Lipofectamine (Invitrogen) was performed according to the manufacturer’s instructions.

For immunoprecipitation, HEK293T cells were transfected with the indicated plasmids when cells reached 60%–80% confluence. 48h post-transfection, cells were lysed with mild lysis buffer (50mM HEPES at pH 7.5, 150mM NaCl, 1mM EDTA, 1% NP-40, 10mM pyrophosphate, 10mM glycerophosphate, 50mM NaF, 1.5mM Na_3_VO_4_, protease inhibitor mixture (Roche), 1mM DTT, 1mM PMSF). Anti-Flag or anti-Myc antibodies were added to the cell lysate and incubated at 4°C for 2h to bind Flag-YAP1 or Myc-TEAD4, and then Pierce Protein A/G agarose beads were added to precipitate the antibody-substrate complex at 4°C for 2h. GFP-Trap agarose beads were directly added to the cell lysis to precipitate GFP-AP-2γ. The immuno-precipitates were washed three times with mild lysis buffer and boiled with SDS sample buffer before analysis by western blot.

#### Quantification and Statistical Analysis

Statistical parameters are reported in the figures and figure legends. Data is considered significant if p < 0.05. Statistical analysis was performed using GraphPad Prism software.

## Supplementary Figure Legends

**Figure S2.**
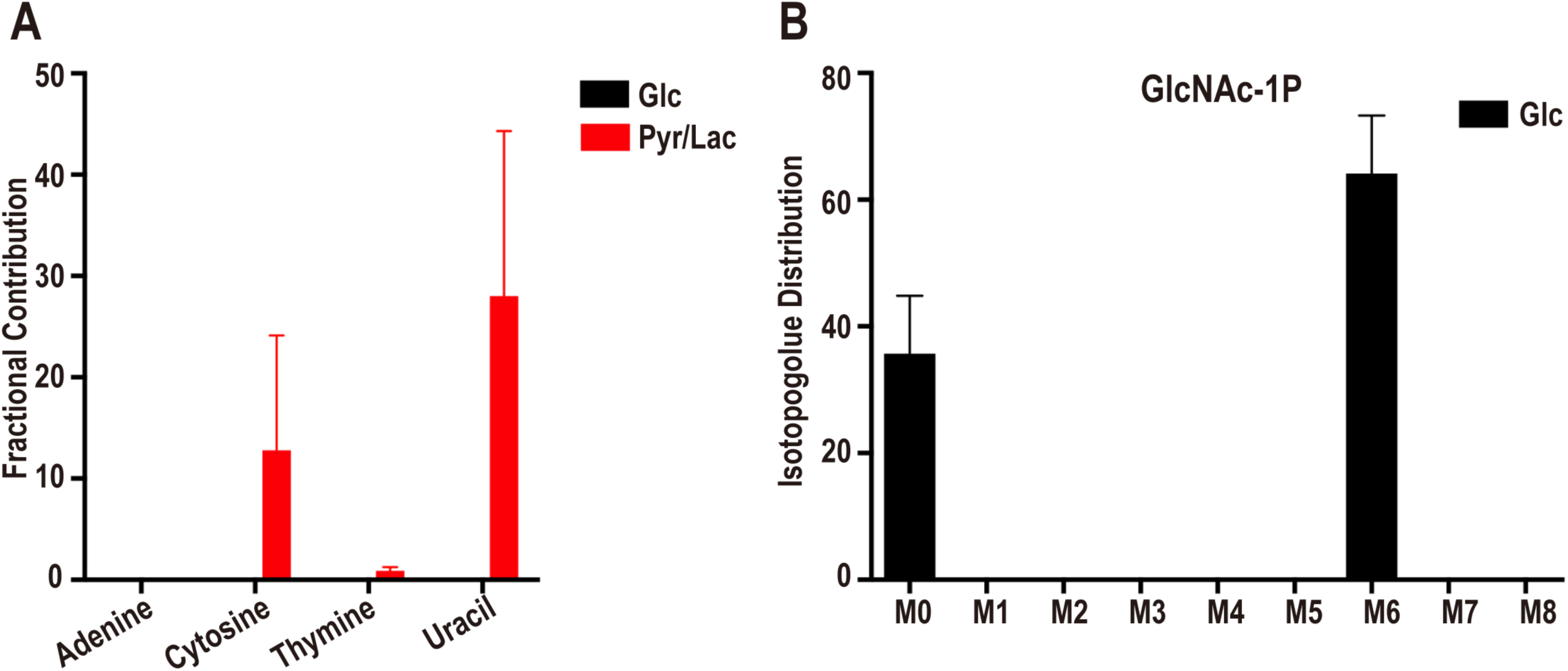
Metabolomic analysis of the compacted morula. (**A**) Contribution of glucose and pyruvate/lactate to nucleobases. U-^13^C-glucose does not contribute carbon to either the purine bases (adenine) or to the pyrimidine bases (cytosine, uracil). In contrast, U-^13^C-pyruvate/lactate contribute to the pyrimidines, but not the purine bases. The nucleobases are generated from the breakdown of nucleotides, and their labeling pattern is an indirect read-out of the labeling of bases within the nucleotides. (**B**) U-^13^C-glucose isotopologue distribution of HBP related metabolite GlcNAc-1P. The most abundant isotopologue is M6 in which 6 of the donated carbons are from exogenous (^13^C) glucose. M0 peak corresponds to unlabeled metabolites from internal resources.

**Figure S3.**
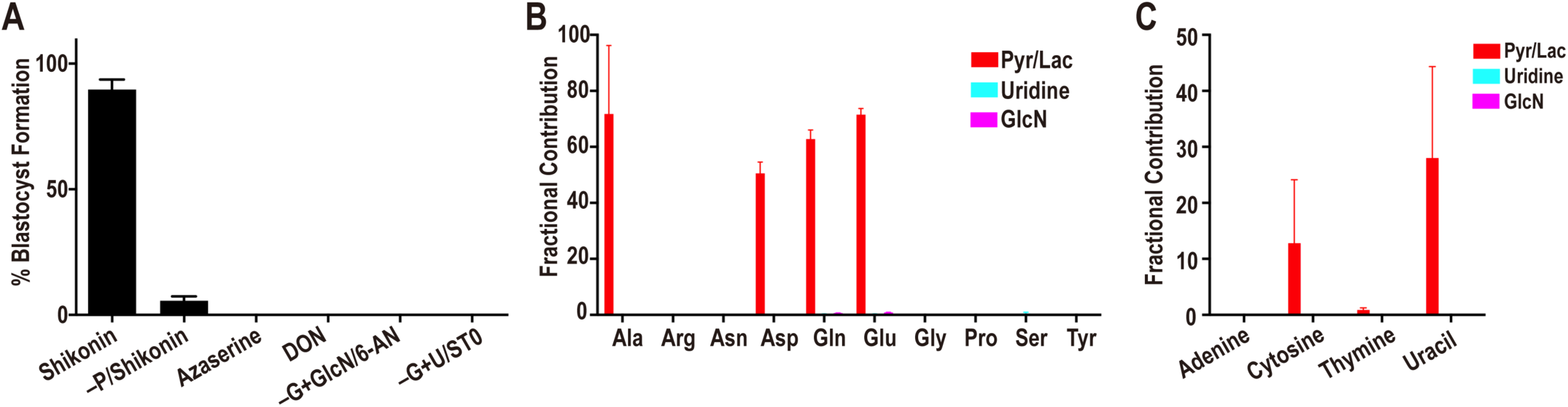
Metabolic contribution of glycolysis, PPP and HBP. (**A**) Percentage of embryos that form blastocysts when cultured with added inhibitors or nutrient supplements. Inhibition of glycolysis (PKM2) by shikonin in +G has no adverse effect on blastocyst formation. However, in the absence of pyruvate (+G–P), shikonin blocks blastocyst formation. Inhibition of HBP by azaserine (2µM), or DON (20µM), both block the morula to blastocyst transition. Embryos cultured in medium containing glucosamine (GlcN) in place of glucose remain sensitive to inhibition of the PPP by 6-AN (2µM). Uridine (U) does not rescue a block caused by inhibition of OGT by ST045849 (ST0, 0.8µM). (**B**) Metabolomic analysis shows that, unlike pyruvate/lactate (red), uridine (light blue) and glucosamine (purple) do not contribute carbons to non-essential amino acids. (**C**) As seen with glucose, labeled uridine and glucosamine do not contribute carbon to nucleobase biosynthesis. In contrast, labeled pyruvate/lactate contribute carbon to pyrimidine base biosynthesis.

**Figure S4.**
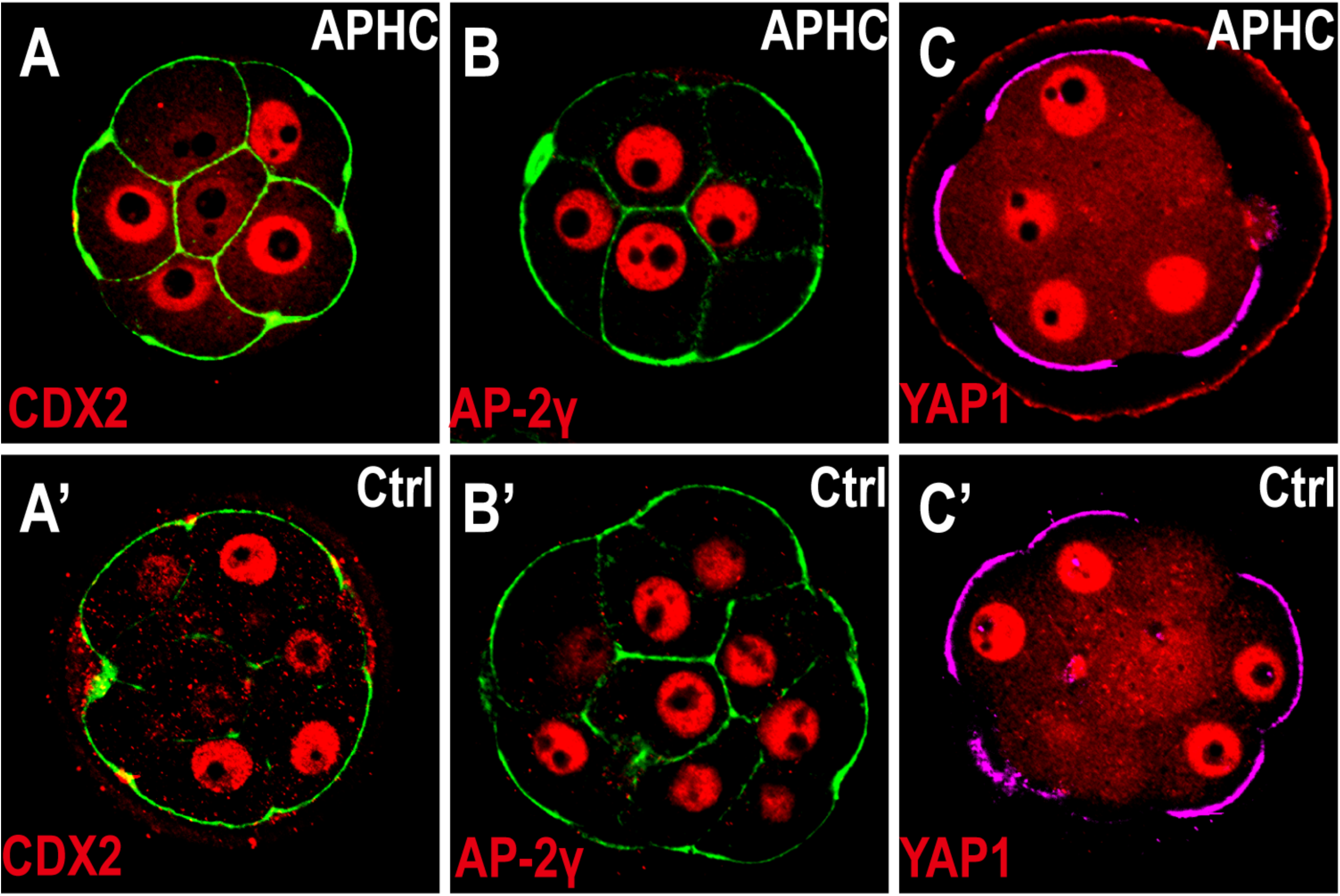
TE fate defect is not caused by 8-cell block. Cell membranes (green) are marked using phalloidin-FITC (**A, A’, B, B’**), and apical surface (purple) are marked by pERM (**C, C’**). Aphidicolin (APHC, 10µM) was added to the embryos at the early 8-cell stage (66h) to inhibit the DNA synthesis and cell cycle progression. (**A, A’**) At 78h control embryos (**A’**) are 8-16 cells and CDX2 (red) is expressed in the polar cells. Aphidicolin treated embryos are arrested at the 8-cell stage (78h), but have high levels of CDX2 expression (**A**). Aphidicolin treated embryos (**B**) have high levels of AP-2γ (red) that is similar to that seen in control embryos (**B’**). (**C, C’**) YAP1 nuclear localization (red) in aphidicolin treated embryos (**C**) is similar to that seen in the polar cells of control embryos (**C’**).

**Figure S5.**
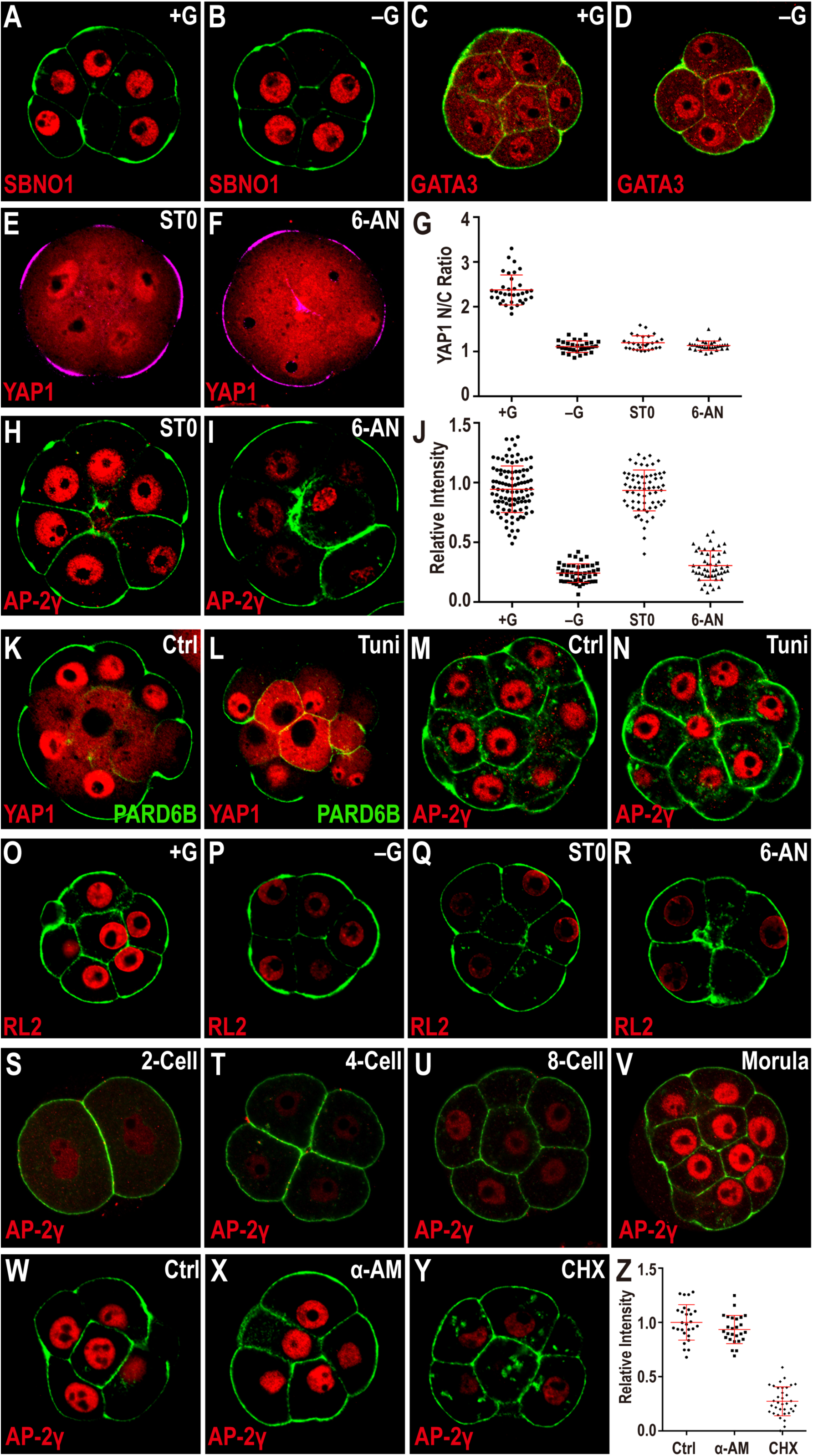
Glucose dependent activation of transcription factors. Cell membranes (green) are marked using phalloidin-FITC, and apical surface are marked by pERM (purple in **E, F**) or PARD6B (green in **K, L**). (**A-D**) Expression of SBNO1 (**A, B**) or GATA3 (**C, D**) in morula stage (78h) embryos is not dependent on exogenously provided glucose. (**E-G**) PPP and HBP and YAP1 nuclear localization. In +G control (see Figure 5A) YAP1 is localized to the nucleus in pERM (purple) marked polar cells. Inhibition of *O*-glycosylation by ST045849 (0.5µM) (**E**) or PPP by 6-AN (2µM) (**F**) both give results similar to that of –G (see Figure 5B). (**G**) Quantitation of YAP1 nuclear to cytoplasmic (N/C) ratio. (**H-J**) In control medium AP-2γ (red) is expressed in the nuclei of all blastomeres (see Figure 5D). This expression is insensitive to *O*-linked glycosylation inhibition by ST045849 (0.5µM) (**H**), but is lost when the PPP is inhibited by 6-AN (2µM) (**I**). Quantitation shown in (**J**). (**K-N**) *N*-linked glycosylation and TE specification. Apical cells with high levels of PARD6B (green) on their membrane also show nuclear localization of YAP1 (red) (**K**). Blocking *N*-glycosylation by tunicamycin (Tuni) treatment causes partial disruption in PARD6B apical localization (**L**). Apical cells that have lost apical PARD6B do not express YAP1 in the nucleus, but in other apical cells that maintain PARD6B expression YAP1 is localized in the nucleus (**L**). Tunicamycin treatment (**N**) does not influence the expression of AP-2γ (**M, N**). (**O-R**) *O*-linked glycosylation is glucose, HBP and PPP dependent. *O*-linked glycosylation is detected using the RL2 antibody and decreases in embryos grown without glucose (**P**), or when *O*-linked glycosylation is inhibited by ST045849 (**Q**), or when the PPP is inhibited using 6-AN (**R**). (**S-V**) AP-2γ expression during preimplantation development. AP-2γ expression is low at the 2-cell (**S**), 4-cell (**T**), and early 8-cell (**U**) stages, and then increases at the compacted morula (**V**) stage. (**W-Z**) AP-2γ expression is sensitive to inhibition of translation. Blocking translation by cycloheximide (CHX) (**Y**), but not transcription inhibition by α-amanitin (α-AM) (**X**) causes a reduction in nuclear AP-2γ levels. (**Z**) Quantitation of nuclear AP-2γ.

**Figure S6.**
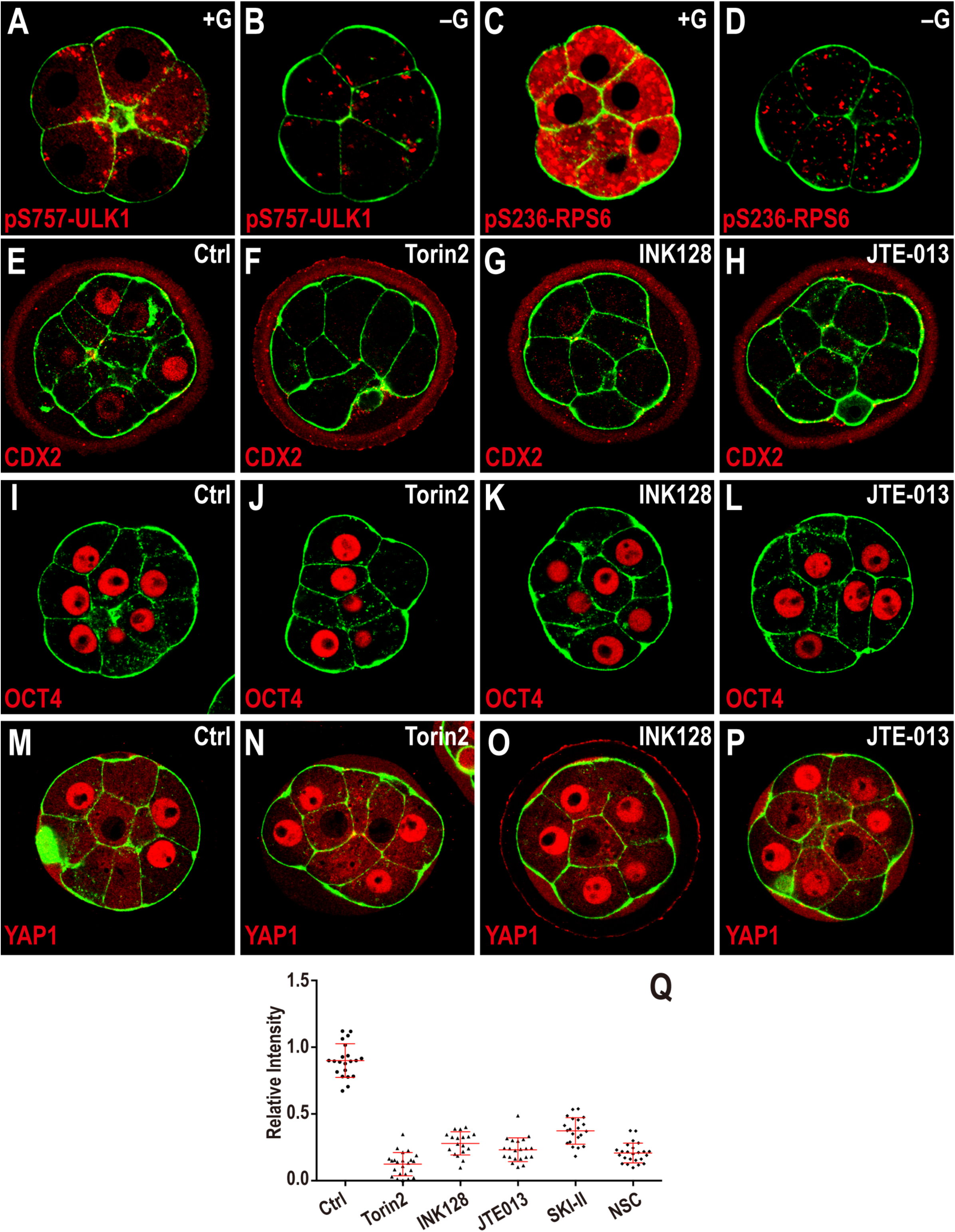
Activation of mTOR by glucose and S1P. Embryos are isolated at 18h and cultured in either +G or –G media until 78h, and inhibitors were added to the medium at 50h. Cell membranes are marked by phalloidin-FITC staining (green). (**A-D**) mTOR target expression requires glucose. In control embryos high levels of pS757-ULK1 (**A**) or pS236-RPS6 (**C**) are seen at 78h. Embryos cultured in –G medium show a dramatic decrease in the levels of pS757-ULK1 (**B**) and pS236-RPS6 (**D**). (**E-H**) CDX2 expression is lost upon mTOR inhibition with Torin2 (**F**) and INK128 (**G**) and inhibition of S1P signaling by JTE-013 (**H**). (**I-P**) Inhibition of mTORC1 and S1P signaling does not perturb OCT4 and YAP1 expression. mTOR inhibition with Torin2 (**J, N**) and INK128 (**K, O**) and S1P signaling by JTE-013 (**L, P**) does not affect OCT4 (**I-L**) and YAP1 (**M-P**). (**Q**) Quantitation of phosphorylation of 4E-BP1 in embryos in which the mTOR and S1P pathways are inhibited (the control data are shared with Figure 6D).

